# Developmental signaling reveals functionally enriched human-specific gene regulation in telencephalic progenitors

**DOI:** 10.64898/2026.07.13.738326

**Authors:** Reed C. McMullen, Bryan J. Pavlovic, Dani M. Swope, David S. Aley, Nathan K. Schaefer, Alex A. Pollen

## Abstract

Comparative transcriptomic studies of neural progenitors implicated in human brain expansion have identified extensive baseline gene expression divergence, yet these differences are weakly enriched for functions relevant to development and disease. This suggests that functionally important regulatory divergence may emerge only under specific developmental signaling conditions. Here, we profiled morphogen-dependent gene expression responses in matched telencephalic neuroepithelial cells (telNECs) from human, chimpanzee, and orangutan at the onset of cortical neurogenesis. Baseline interspecies differences were extensive but functionally diffuse. In contrast, a distinct set of genes exhibited species-divergent responses to morphogen stimulation despite conserved baseline expression. These response genes were strongly enriched for regulators of progenitor proliferation and differentiation, neurodevelopmental disorder risk genes, and loci harboring human-lineage sequence changes. Together, these findings show that developmental signaling exposes a functionally enriched class of regulatory divergence beyond baseline comparisons and provide a framework for identifying evolutionarily relevant gene regulation during human brain development.

## INTRODUCTION

The three-fold expansion of the human brain since our divergence from a common ancestor with chimpanzees represents one of the most conspicuous human traits. By mid-gestation the human brain is already twice the size of the chimpanzee brain,^1^ suggesting that species differences emerge at early stages of prenatal development. The radial unit hypothesis proposes that increased proliferation of telencephalic neuroepithelial cells (telNECs) during early brain expansion increases the number of radial glia cells, which produce an increased number of neurons, expanding cortical surface area.^2,3^ Consistent with this model, initial studies suggest that human telNECs exhibit increased proliferation,^4^ prolonged metaphase,^5,6^ and altered NEC morphology.^7^ However, comparisons of human and chimpanzee pluripotent stem cell (PSC)-derived organoids have largely focused on later stages of neurogenesis, revealing divergence in radial glia gene expression and signaling pathway activation,^8,9^ rather than examining earlier stages of NEC expansion when proliferative differences that drive brain size first emerge.^7^

Gene expression divergence is thought to underlie evolved differences in cell behavior.^10–12^ However, most gene expression divergence reflects neutral drift rather than functional divergence,^13–15^ consistent with patterns at the genome sequence level.^16^ In line with this model, baseline gene expression variation shows poor overlap with the genetic variants linked to complex traits in humans, including neuroanatomical phenotypes.^17,18^ Despite this limited overlap, studies of human-specific transcriptome evolution have largely relied on steady-state comparisons of gene expression.^8,9,19–21^ Together, these observations suggest that baseline gene expression comparisons may be poorly suited to prioritize the subset of regulatory changes that are functionally relevant for brain expansion.

One approach to identifying trait-linked variation that is missed in baseline contexts is to focus on genomic responses to physiological stimuli.^17,18,22,23^ Adaptive regulatory variants that alter gene expression only in specific cellular, environmental, or developmental contexts can influence phenotypes while minimizing deleterious effects on baseline gene expression programs. Recent studies underscore the potential of physiological stimuli to link trait-associated variants to gene regulatory variation,^24–32^ while enriching for genetic variants under adaptive selection.^17,33–35^ Although these studies have largely focused on genetic variation among humans, comparing stimulus-dependent responses between species recently uncovered functionally relevant divergence in cardiomyocytes.^36^ However, the evolution of stimulus-dependent responses remains largely unexplored in the hominid lineage,^23^ particularly in the nervous system.

During nervous system development, morphogens and mitogens play central roles in regulating NEC patterning, proliferation, and differentiation.^37–39^ Responses to these signals thus represent a potential substrate for selection on brain size and structure. Several pathways have been implicated in cortical expansion based on functional experiments, genetic changes, and gene expression divergence, including Wnt,^17,40–42^ Notch^43–45^, mTOR^8,46,47^, and retinoic acid^48,49^ signaling. More recently, FGF-ERK-BMP7 signaling in radial glia was proposed to contribute to mammalian cortical expansion.^50^ Despite the strong effects of morphogens on cortical development, whether human and chimpanzee differ in their transcriptional responses to these pathways during NEC expansion—and whether such differences contribute to human-specific developmental programs— remains unknown.

We predicted that systematic morphogen perturbation of telNECs would expose functionally relevant human-specific gene expression divergence. To test this hypothesis, we combined a phylogeny-in-a-dish approach^51^ with multiplexed morphogen perturbations,^52–55^ pooling PSC-derived telNECs from human, chimpanzee, and orangutan and measuring single cell responses to morphogens along a developmental timecourse.^56^ Directed differentiation of PSCs provides a scalable reductionist model system that recapitulates key aspects of developmental patterning, timing, cell type specification, and interspecies differences^4,8,57,58^. By modulating nine signaling pathways active during telNEC expansion using 23 distinct activators and inhibitors, we discovered conserved and human-specific stimulus-dependent gene regulatory changes and assigned divergence to underlying signaling mechanisms. Among stimuli, inhibition of Notch and activation of FGF and BMP signaling elicited the strongest transcriptional and cellular effects. In contrast to baseline gene expression differences, stimulus-dependent responses were enriched for genes related to brain development and patterning. Human-specific responses to FGF and BMP7 stimulation were further enriched for genes regulating cell cycle, brain size, and neuropsychiatric disorders, as well as loci harboring human-specific sequence variation. Together, these findings demonstrate that morphogen perturbation can systematically reveal functional regulatory divergence masked in baseline gene expression comparisons and provide a framework for dissecting evolved developmental signaling responses across hominids.

## RESULTS

### A multiplexed morphogen screen of interspecies great ape telencephalic cultures

We characterized morphogen responses in telNECs during the expansion phase preceding cortical neurogenesis across great ape species (Fig. 1A-C), when telNECs *in vitro* retain regional plasticity,^59^ produce neurons,^58,60,61^ and respond to patterning signals. To distinguish interspecies divergence from interindividual variation,^62^ we differentiated PSC lines from six human, six chimpanzee, and one orangutan individual to telNEC fate (Fig. S1A, Table S1, Methods),^63–65^ balancing species and sex across four differentiation batches.

**Fig. 1.**
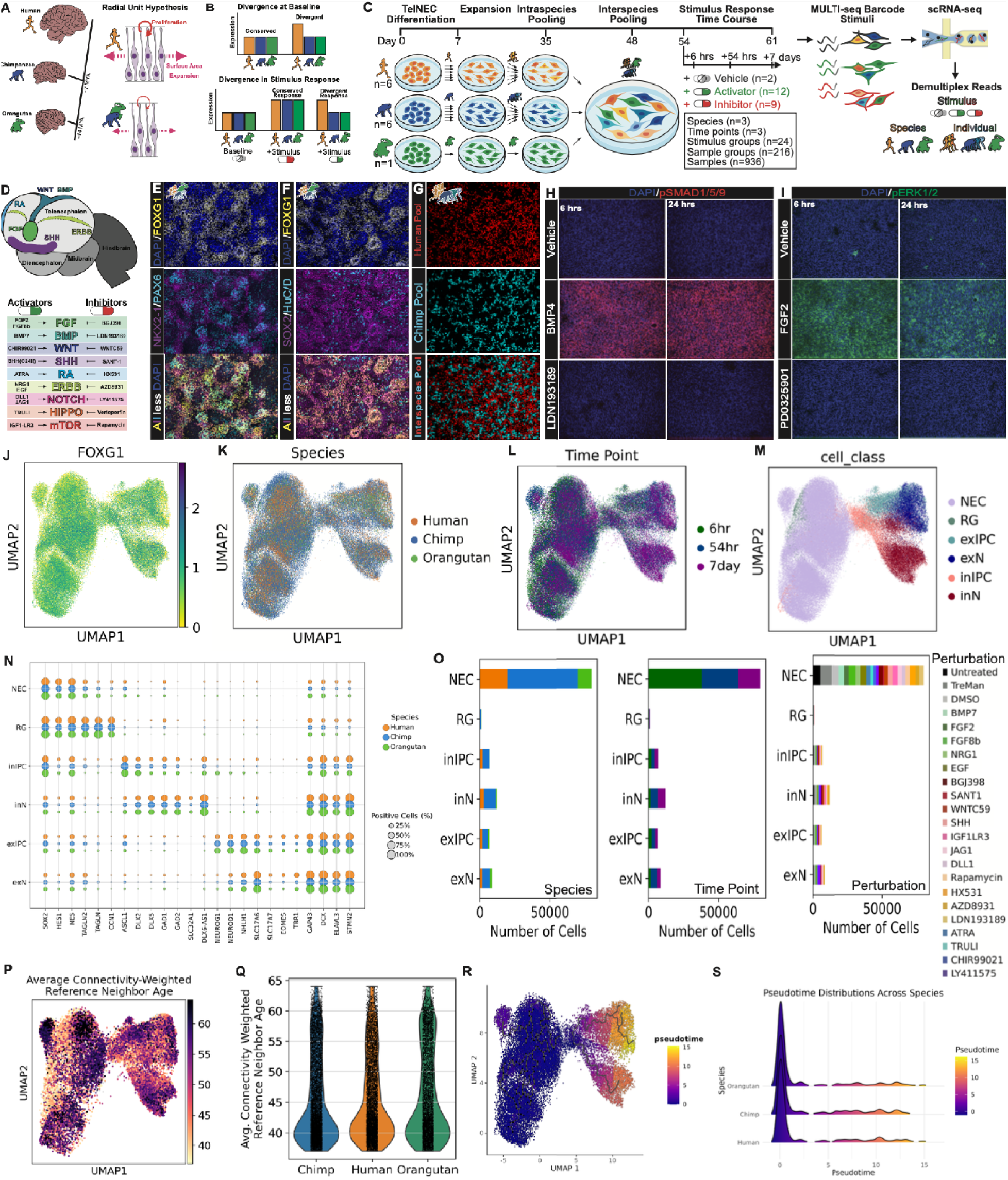
A multiplexed morphogen screen of mixed species great ape forebrain cultures. **(A)** Schematic of great ape phylogeny with approximate divergence times and relative adult brain sizes, alongside the radial unit hypothesis. MYA, million years ago. **(B)** Conceptual framework distinguishing gene expression divergence at baseline versus stimulus-dependent responses, highlighting that some regulatory differences may be undetectable at baseline but emerge under stimulation. **(C)** Experimental design. PSC-derived telencephalic neuroepithelial cells (telNECs) from human, chimpanzee, and orangutan were differentiated, pooled within and across species, and exposed to activators or inhibitors of nine developmental signaling pathways. Cells were profiled by multiplexed scRNA-seq across three time points (6 h, 54 h, 7 d), followed by demultiplexing by species, individual, and perturbation. **(D)** Schematic of forebrain patterning centers and associated morphogen pathways, adapted from Montiel and Aboitiz 2015.^120^ Dashed lines indicate caudal telencephalic position. Table lists pathway activators and inhibitors used. **(E–G)** Immunocytochemistry (ICC) validation of telencephalic identity and species mixing. (E–F) Marker expression (FOXG1, PAX6, NKX2-1, SOX2, ELAVL3/4) in interspecies cultures at day 54. (G) CellTrace labeling confirms mixing of intraspecies pools prior to interspecies pooling. **(H–I)** ICC validation of pathway activation starting at day 21. (H) BMP pathway activation assessed by pSMAD1/5/9 following BMP4 (25 ng/mL) treatment and inhibition by LDN193189 (0.5 µM). (I) FGF pathway activation assessed by pERK1/2 following FGF2 (50 ng/mL) treatment and inhibition by PD0325901 (0.5 µM). **(J–M)** UMAP embedding of 114,711 FOXG1+ telencephalic cells colored by (J) FOXG1 expression, (K) species, (L) time point, and (M) cell class. **(N)** Dot plot of canonical marker gene expression across cell classes, split by species; dot size indicates fraction of expressing cells. **(O)** Composition of cell classes across species, time points, and perturbations. **(P–Q)** Mapping to a rhesus macaque reference atlas. UMAP colored by connectivity-weighted reference age (P) and corresponding distributions by species (Q). **(R–S)** Pseudotime analysis. UMAP colored by pseudotime (R) and distributions across species (S), showing comparable developmental progression.

We expanded telNECs in media optimized for cortical progenitor maintenance^60^ (Fig. S1A, Table S1) and pooled individuals first within species (intraspecies) and then across species (interspecies, Fig. 1C), a strategy that stabilizes growth, controls for cell-extrinsic effects, and mitigates batch effects.^51,66–68^ A majority of human,chimpanzee, and orangutan cells were of FOXG1+ telencephalic fate, with balanced ventral (NKX2-1+) and dorsal (PAX6+) identities both prior to (∼1:1 ratio, Fig. S1B-C) and following pooling, with 15.0% of cells representing telencephalic IPCs or neurons (FOXG1+/HuC/D+, Fig. 1F, Fig. S1D-E), indicating that cultures were predominantly telencephalic progenitors with capacity for neurogenesis.

To select signaling pathways, we screened morphogens implicated in telencephalon development,^40–42,47,48,50^ diseases affecting brain size,^69,70^ and interindividual variation in neuroanatomical traits^71–76^ (Fig. 1H-I, Fig. S1F-M, Table S2). We selected 21 ligands or small molecules activating or inhibiting 9 signaling pathways implicated in brain development: BMP, FGF, ERBB, WNT, mTOR, NOTCH, RA, HIPPO, and SHH (Fig. 1B).

To quantify species divergence in morphogen response with single cell resolution, we treated interspecies cultures with activators or inhibitors at an acute stage (6 hours), near the first cell division (54 hours), or after extended differentiation when cell type composition differences might emerge (7 days) (Fig. 1C). Prior to these treatments, we removed media factors supporting continued expansion for six days to focus on the effects of treatments. Cells were labeled with MULTI-seq barcodes and profiled by multiplexed scRNA-seq. Following sequencing and alignment (Fig. S2E-I), we performed species and individual demultiplexing, perturbation assignment, doublet detection, and ambient RNA removal using CellBouncer^77^ (Fig. S2A-D), followed by standard preprocessing and quality control.^78^ We captured 207 of 216 (95.8%) expected sample groups (defined as each combination of species, perturbation, and time point) and 892 of 936 (95.3%) expected samples (defined as each combination of sample group and individual)

Among 114,711 *FOXG1*+ telencephalic cells (Fig. 1J, Fig. S2L-O, S3), unsupervised clustering accounting for biological and technical variation^79^ revealed homologous cell types across species and timepoints (Fig. 1K-L, Methods). Cells were predominantly telNECs (69.6% average), with intermediate progenitor cell (IPC) and immature neuron (immN) classes (Fig. 1M-N, Fig. S3A, Table S3, Methods). Subsetting and integration^80^ demonstrated both excitatory and inhibitory lineages, including neurons with preplate,^81–83^ ventral, ventromedial, and GnRH neuron identities, with comparable regionalization across species (Fig. 1M-N, Fig. S3B, Fig. S4D-F, Table S3, Methods). Across all samples, TelNEC abundance decreased from 82.8% to 55.5% from 6 hrs to 7 days while immNs increased from 6.8% to 28.5%, indicating active neurogenesis, and consistent with effects differentiation in vehicle-treated cells (Fig. S3E). Perturbations were evenly represented across species, time points, and cell classes (Fig. 1O, Fig. S3).

To assess the homology of telNECs across species and developmental stage, we mapped our dataset to a reference atlas of developing rhesus macaque telencephalon,^84^ which is equally divergent from human and chimpanzee. Using semi-supervised probabilistic annotation with scANVI,^85^ telNECs from all species mapped to comparable first trimester stages (mean macaque post conception day: 43.0 human, 43.5 chimpanzee, and 43.9 orangutan) (Fig. 1Q). Pseudotime analysis,^86^ indicated similar distributions of cells along a neurogenic trajectory across species (Fig. 1R-S). Together, these results establish that pooled interspecies cultures generate comparable telencephalic progenitor and early neuronal populations across species.

### BMP7 and FGF activation and Notch inhibition elicit the strongest responses in telNECs

To characterize morphogen responses across stimuli and species (Fig. 2A), we first confirmed that each stimulus elicited the expected pathway activity, with activator and inhibitor pairs demonstrating reciprocal regulation of canonical effector genes (Fig. 2B, Table S4). We next quantified response strength by comparing each condition to its corresponding vehicle control using two complementary measures: the number of differentially expressed genes (response genes, rGenes; FDR < 0.10) from pseudobulk linear mixed models^87^ and the energy distance (E-distance),^88^ which captures shifts in both gene expression and cell state composition. Both metrics were independent of cell number (Fig. S5A–H), and provided concordant measures of response strength (Spearman’s rho = 0.85, p = 2.7e-30; Fig. S5I, Table S5).

**Fig. 2.**
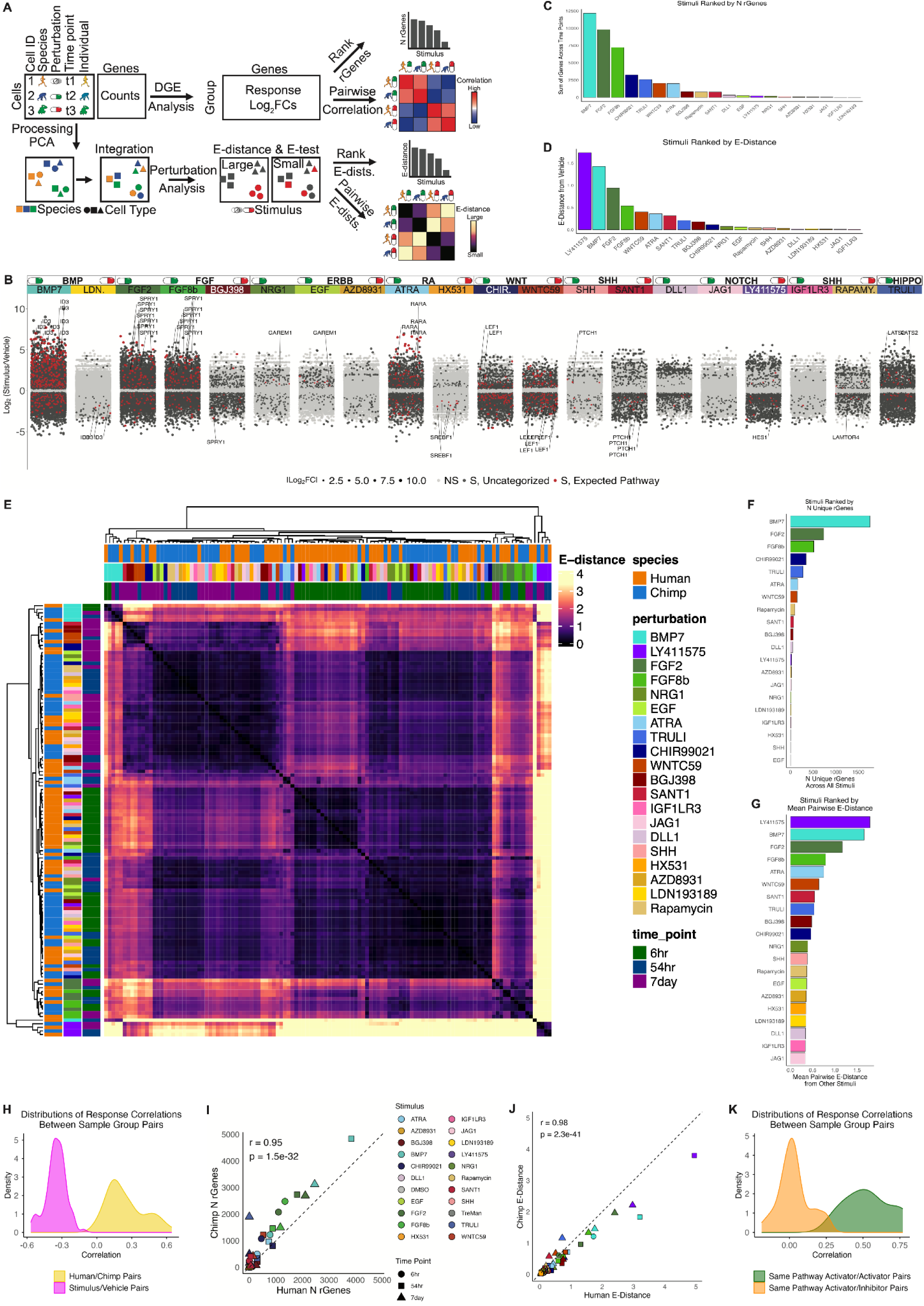
BMP and FGF activation and Notch inhibition elicit the largest effect on telNECs. **(A)** Overview of analysis workflow. Single-cell data were processed to quantify morphogen responses using differential gene expression (rGenes) and energy distance (E-distance), enabling ranking of stimuli by response magnitude and comparison of response distinctiveness across conditions. **(B)** Gene-level responses across stimuli. Log₂ fold changes (stimulus vs vehicle) are shown for each perturbation, with genes colored by significance (FDR < 0.10) and annotated by canonical pathway effectors. Activator–inhibitor pairs show reciprocal regulation of pathway targets. **(C–D)** Ranking of stimuli by response magnitude. Stimuli are ordered by total number of rGenes (C) and E-distance (D), identifying BMP7, FGF2/8b, and Notch inhibition (LY411575) as the strongest perturbations. **(E)** Heatmap of pairwise E-distance across sample groups, revealing clustering by stimulus and time point rather than species for strong stimuli. **(F–G)** Distinctiveness of responses. Stimuli are ranked by the number of unique rGenes (F) and mean pairwise E-distance (G), indicating that strong-effect pathways also produce the most distinct transcriptional responses. **(H)** Distribution of gene expression correlations between species-matched samples (human vs. chimpanzee) and perturbation-matched samples (stimulus vs. vehicle) for perturbations with more than 50 rGenes, showing stronger divergence across perturbations than across species. **(I–J)** Cross-species concordance of response magnitude. The number of rGenes (I) and E-distance (J) are highly correlated between human and chimpanzee across conditions. **(K)** Correlation structure across pathway perturbations. Responses are positively correlated within pathways (activator–activator) and inversely correlated between activators and inhibitors of the same pathway.

Across conditions, BMP7 and FGF signaling (FGF2 and FGF8b) elicited the strongest responses by both rGene count and E-distance, while the Notch inhibitor LY411575 showed a comparably large effect by E-distance (Fig. 2C–G). Gene expression responses peaked at 54 hours, whereas E-distances continued to increase through 7 days, consistent with accumulating changes in cell state composition (Fig. S5K–O).

Stimuli targeting the same pathway produced correlated gene expression responses, while activator–inhibitor pairs were anticorrelated (Fig. 2K, Table S6). Among the strongest stimuli, FGF2 and FGF8b showed similar responses and clustered with ERBB pathway activators (NRG1, EGF), while BMP7 shared substantial overlap in response genes with the WNT activator CHIR99021 (Fig. 2E, Fig. S5Q–S). Both response strength and distinctiveness increased over time (Spearman’s ρ = 0.70).

Morphogen responses were highly conserved across species. Human and chimpanzee rGene counts and E-distances were strongly correlated (Pearson’s r = 0.95 and 0.98, respectively; Fig. 2I–J), and gene expression responses in matched sample groups were similarly correlated across species (Fig. 2H, Table S6). For strong-effect stimuli, samples clustered primarily by stimulus and timepoint rather than species (Fig. 2E). Together, these analyses identify a subset of developmental signaling pathways that dominate transcriptional responses in telNECs and are highly concordant across species, motivating further examination of their impact on cell state and fate.

### Cell state and fate responses to morphogens are conserved across great ape species

Beyond transcriptional responses, morphogen stimulation also reshaped cellular composition. Two transcriptomic clusters were almost entirely defined by stimuli: one marked by canonical BMP response genes (*SMAD6/9*, *BAMBI*, and *NOG)* and another by FGF/ERBB response genes (*DUSP4*/6, *SPRY1*/*4)* (Fig. 3A-B, Fig. S6A-G, Table S3).

**Fig. 3.**
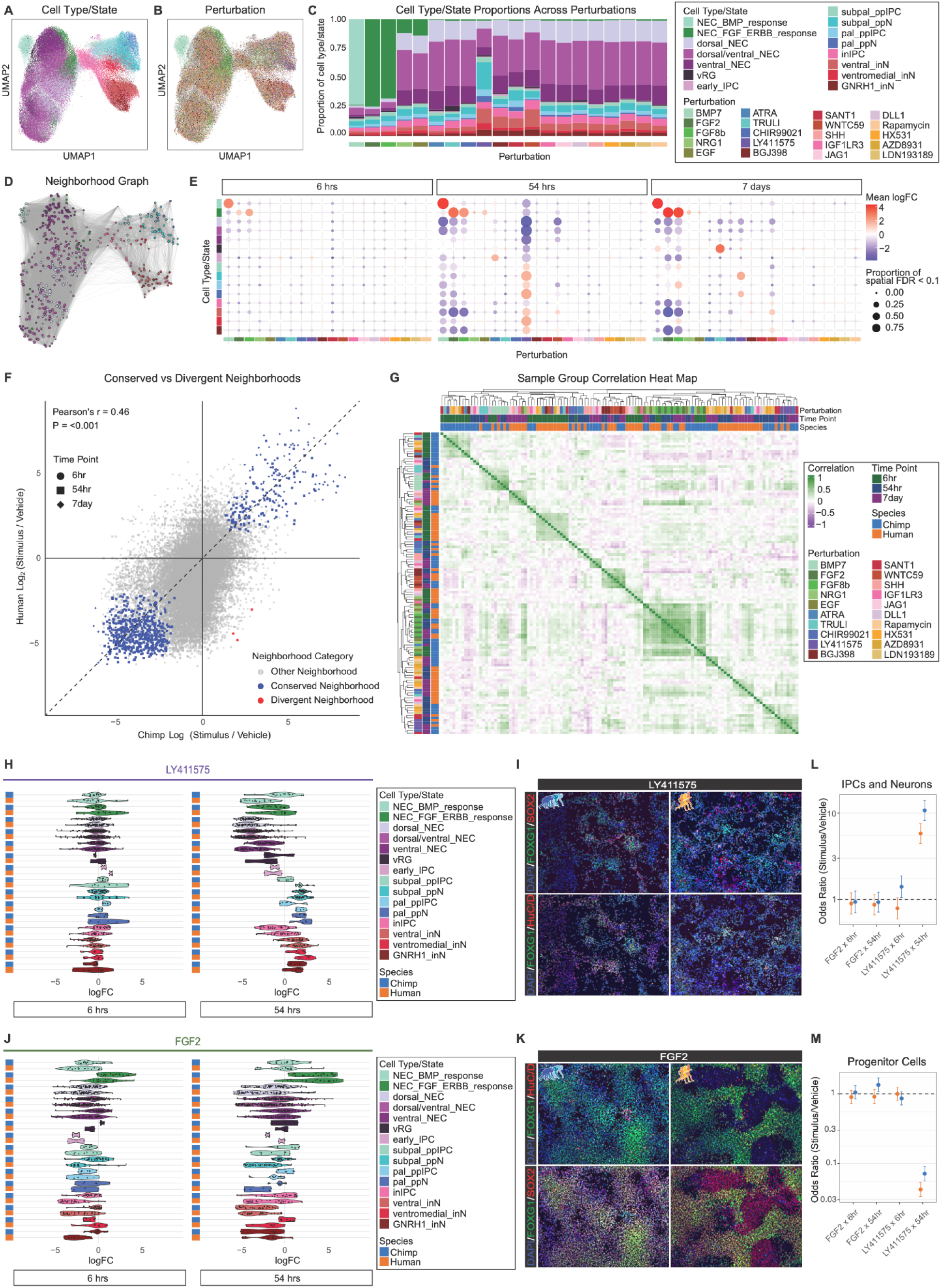
Morphogen-induced cell state and fate responses are largely conserved across species. (A–B) UMAP embedding of telencephalic cells colored by (A) cell type/state annotation and (B) perturbation, showing broad overlap of cell states across conditions. **(C)** Cell type/state composition across perturbations, indicating stimulus-dependent shifts in cellular abundance. **(D)** Graph representing fine-grained cellular neighborhoods. Nodes correspond to neighborhoods (sized by cell number) and are colored by majority cell type/state identity; edges reflect shared cellular membership. **(E)** Differential abundance of cell states across perturbations and time points. Dot size indicates the fraction of neighborhoods within a cell type/state with significant changes (FDR < 0.10), and color indicates mean log₂ fold change. **(F)** Cross-species comparison of stimulus-induced abundance changes. Most significant responses are conserved between human and chimpanzee (blue), with few divergent neighborhoods (red). **(G)** Hierarchical clustering of pairwise Pearson’s correlation coefficients. For perturbations with strong abundance effects, samples cluster by perturbation identity rather than by species or timepoint, consistent with broad cross-species conservation. **(H-M)** Cell state–specific abundance responses and ICC validation to Notch inhibition (LY411575) (H) and FGF2 stimulation (J) across time points in human and chimpanzee. ICC validation of LY411575 (I) and FGF2 (K) responses in intraspecies cultures, with quantification showing changes in IPC/neuron abundance (L; DAPI⁺/FOXG1⁺/ELAVL3/4⁺) and progenitor abundance (M; DAPI⁺/FOXG1⁺/ELAVL3/4⁻/SOX2⁺) across treatments and time points.

We next used a cluster-free differential abundance method to quantify fine-grained shifts in cellular neighborhood composition across developmental and stimulus-response axes^89^ (Fig. 3D–E, Table S7). Notch inhibition with LY411575 at 54 hrs elicited the strongest abundance changes, followed by BMP7, FGF2/8b, and WNTC59 (Fig. 3E). Perturbations targeting the same pathway produced concordant abundance changes across timepoints (Fig. 3C,E), indicating coherent pathway-level effects. While LY411575 and WNTC59 drove delayed increases in neuronal abundance (Fig. 3E,H), BMP7 and FGF2/8b rapidly increased distinct telNEC neighborhoods as early as 6 hrs (Fig. 3E,J, Fig. S6D,F). The rapid onset of these BMP and FGF effects, in the absence of differentiation markers, is consistent with an initial shift in cell state rather than cell fate, indicating that early morphogen responses reconfigure progenitor states before altering lineage output.

Cell fate responses were broadly consistent with known developmental trajectories. WNTC59 decreased preplate neuron abundance while increasing inhibitory IPCs and neurons, consistent with the dorsal origin of preplate neurons and ventral origin of inhibitory neurons (inNs) (Fig. 3E, Fig. S6C). However, WNT activation with CHIR99021 did not reciprocally increase preplate neuron abundance (Fig. 3E), and SHH perturbation had no significant effect on neuronal lineage abundance (Fig. 3E). BMP7 reduced InN abundance at 7 days while sparing preplate neurons and inducing pallial (HEY1, ROR2) and cortical hem (MSX1, LMO2, ID1/2/3, DOK5) markers in NECs, consistent with BMP7 localization during development (Fig. 3E, Fig. S6D).^84,90,91^

These cellular responses were broadly conserved across species. Abundance changes between human and chimpanzee were positively correlated across neighborhoods (Pearson’s r = 0.46, *p* < 0.001; Fig. 3F), strong-effect stimuli clustered by stimulus and time point rather than species (Fig. 3G), and only 3 of 514 neighborhoods showed significant species differences, all within the BMP7 response at 54 hrs.

For orthogonal validation, we performed immunocytochemistry (ICC) on intraspecies, multi-individual cultures of human and chimpanzee cultures treated with LY411575 or FGF2 for 6 and 54 hrs. LY411575 decreased the abundance of telencephalic progenitors (FOXG1+/SOX2+/ELAVL3/4−) and increased the abundance of neurons (FOXG1+/ELAVL3/4+) after 54 hrs in both species, with similar effect sizes. In contrast, FGF2 increased culture density without substantially altering cell type composition in either species (Fig. 3J–L).

Together, these results demonstrate that morphogen-induced cell state and fate transitions are largely conserved across great ape species, even for pathways that elicit strong transcriptional responses.

### Morphogen stimuli elicit conserved responses functionally relevant to brain expansion

To distinguish shared from species-specific morphogen responses, we classified response genes (rGenes) based on concordance of stimulus-induced gene expression changes between human and chimpanzee (FDR < 0.10, Methods). Across 928,619 gene-by-stimulus combinations, we identified 41,861 rGenes (4.3%) (Fig. 4A), including 10,047 conserved (23.9%), 1,222 divergent (2.9%), and 30,683 other rGenes (73.1%), which respond significantly in one species only and for which the interaction of species and perturbation is not significant (Fig. 4B-E).

**Fig. 4.**
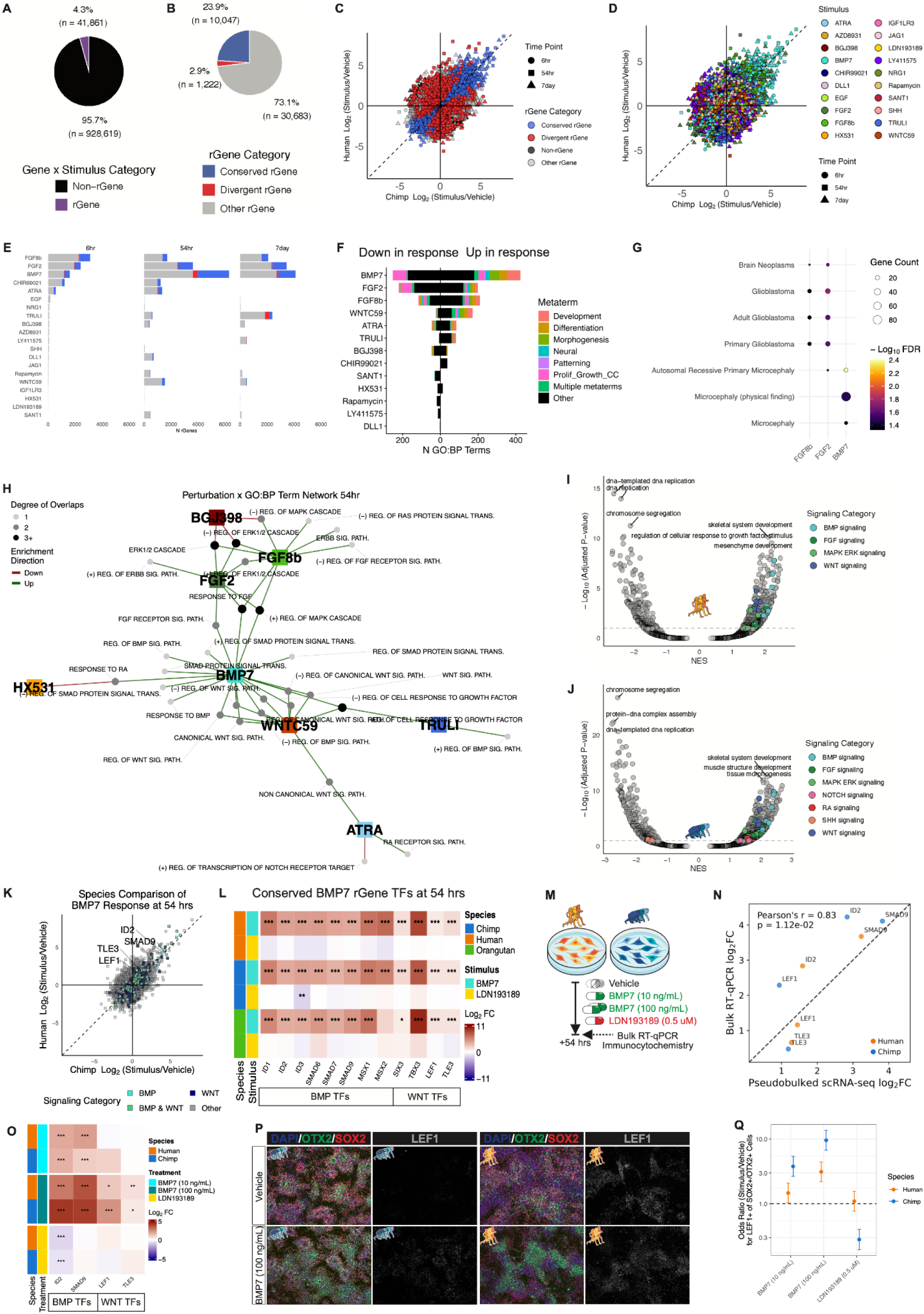
Morphogen stimulation reveals functionally enriched conserved gene responses linked to brain expansion. (A–B) Classification of gene expression responses across all gene–stimulus interactions. A subset of genes show significant stimulus responses (‘rGenes’; FDR < 0.10), the majority of which are conserved between species, with a smaller fraction exhibiting species-divergent responses. **(C–D)** Comparison of human and chimpanzee transcriptional responses across all genes (C) and colored by perturbation (D), showing strong concordance in response magnitude and direction across species. **(E)** Number of rGenes per stimulus across time points, highlighting that BMP and FGF pathway perturbations elicit the largest transcriptional responses. **(F)** Functional enrichment of stimulus-responsive genes. rGenes are enriched for biological processes related to development, proliferation, and differentiation. Number of brain-relevant and other GO:BP terms plotted. **(G)** Enrichment of genes associated with disorders of brain development and proliferation, including microcephaly and glioblastoma, among BMP- and FGF-responsive genes. **(H)** Network representation of enriched GO:BP terms across stimuli at 54 hrs among up- (red) and down- (green) regulated stimulus-response genes, revealing coordinated regulation within and across signaling pathways, including possible feedback regulation of pathway activity (red and green). **(I–J)** Gene set enrichment analysis (GSEA) of BMP7 responses in human (I) and chimpanzee (J), showing conserved enrichment of signaling and cell cycle–related programs. **(K)** Direct comparison of BMP7-induced gene expression changes between species at 54 hrs, highlighting strong conservation of pathway responses. **(L)** Conserved TF responses to BMP7 and its inhibition (LDN193189) across human, chimpanzee, and orangutan, including BMP and WNT pathway effectors. **(M)** Design for validation of conserved BMP7 rGenes at 54 hrs by bulk RT–qPCR and ICC. **(N)** Concordance between pseudobulk scRNA-seq and bulk RT–qPCR measurements of conserved BMP7 responses. **(O)** Gene expression responses to BMP7 pathway modulation by bulk RT-qPCR across conserved rGenes. **(P–Q)** ICC and quantification of LEF1-positive progenitors following BMP7 treatment and inhibition, confirming conserved LEF1 induction at the protein level.

We predicted that conserved rGenes would be enriched for effectors of signaling pathways related to brain expansion. To assess functional relevance, we grouped enriched GO:BP terms into metaterms related to development, patterning, differentiation, proliferation/cell cycle, and neural processes. Whereas non-rGenes showed no GO:BP enrichment, 15 of 20 conserved stimulus-response gene sets were enriched for at least one brain-expansion metaterm (Fig. 4F, Table S8).

Conserved rGenes were enriched for disorders of proliferation, including neoplasms, hyperplasias, and cancers (Table S8). Notably, FGF8b and FGF2 responses were enriched for brain-specific proliferation disorders, including brain neoplasms (FGF8b FDR = 6.1e-3, FGF2 FDR = 7.7e-3) and glioblastoma (FGF8b FDR = 4.4e-2, FGF2 FDR = 1.7e-2), while BMP7 responses were enriched for microcephaly (FDR = 3.5e-2) and autosomal recessive primary microcephaly (FDR = 2.5e-4) (Fig. 4G). Conserved responses to FGF8b and BMP7 were further enriched for neurodevelopmental disorders (NDDs), including autism spectrum disorder, bipolar disorder, schizophrenia, and intellectual disability.

We next examined the dynamics of conserved responses, focusing on FGF2/8b and BMP7 due to their large effect sizes and enrichment for proliferation-related processes (Fig. 4F). At 6 hrs, FGF2 and FGF8b upregulated genes were enriched for canonical FGF–MAPK–ERK signaling, with reciprocal downregulation upon inhibition (BGJ398) (Fig. S6D,J,K). Transcription factors of the ETV (*ETV1/4/5*) and ETS (*ELK3*) families were consistently induced, accompanied by increased expression of progenitor-associated regulators (e.g., *HEY1*, *MYC*) and decreased expression of IPC-associated factors (e.g., *ASCL1*, *OLIG1*), consistent with maintenance of progenitor identity, contrasting with Notch inhibition (LY411575), which upregulated *ASCL1* (Fig. S6F). By 54 hrs, cell cycle-associated genes were unexpectedly downregulated following FGF activation, potentially reflecting density-dependent feedback on proliferation (Fig. S6J,K).

The conserved BMP7 response showed rapid induction of canonical BMP TFs (*SMAD6/7/9*, *MSX1/2*) by 6 hrs, followed by broader activation of BMP pathway genes at 54 hrs, including *BMPR2*, *BMPR1A*, and *BMP7* itself, consistent with positive feedback (Fig. S6B,D,G). BMP7-downregulated genes were enriched for nervous system differentiation (Fig. S7B). As with FGF activation, BMP7 increased progenitor-associated TFs (*HMGA2*, *HEY2*) and decreased IPC-associated TF *ASCL1* (Fig. S6F), consistent with maintenance of progenitor identity. Despite this, cell cycle-associated genes, including MCPH-linked genes, were downregulated at later time points (Fig. 4I,J, Fig. S6I), suggesting feedback regulation of proliferation.

BMP7 responses engage in crosstalk with multiple signaling pathways, including enrichment for FGF–MAPK/ERK signaling and substantial overlap in rGenes with FGF perturbations (Fig. 4H–J, Fig. S6J). In addition to this convergence with FGF signaling, BMP7 induced a second axis of crosstalk with WNT signaling, including conserved upregulation of WNT response TFs (*LEF1*, *TLE3*) and downstream developmental regulators (*SIX3*, *TBX3*) from 6 hrs through 7 days (Fig. 4H, Fig. S6D,E). Consistent with canonical WNT signaling, *LEF1* was inversely regulated by WNT inhibition (WNTC59) and activation (CHIR99021), and BMP7-dependent induction of WNT targets was confirmed by comparison to BMP inhibition (LDN193189) across species (Fig. S6F–H).

We validated these findings by RT-qPCR and ICC in independent human and chimpanzee intraspecies cultures treated with vehicle, BMP7, or BMP inhibitor LDN193189 (Fig. 4M). Expression of BMP-responsive TFs (*ID2*, *SMAD9*) and WNT pathway factors (*LEF1*, *TLE3*) was concordant between single-cell and bulk measurements and showed conserved, dose-dependent induction across species (Fig. 4L,N). Consistent with transcript-level changes, LEF1 protein increased with BMP7 treatment and decreased or remained unchanged with BMP inhibition (Fig. 4O,P; Fig. S6L), supporting BMP–WNT pathway interaction in telNECs.

Together, these results show that conserved morphogen responses define shared, stimulus-dependent developmental programs governing progenitor proliferation, differentiation, and signaling.

### Signaling mechanisms underlying species divergence in gene expression

Having defined a shared, stimulus-dependent regulatory program, we next asked how species divergence emerges within this conserved signaling context by distinguishing baseline differences in gene expression from stimulus-dependent differences in regulatory response (Fig. 5A-C). Baseline divergence between human and chimpanzee telencephalic progenitors was widespread but functionally diffuse. Of 7,677 differentially expressed genes (44.2%, FDR < 0.10), enriched terms primarily reflected housekeeping processes, and the most different genes were largely non-coding (Fig. S8A-C). Only 2.8% of enriched GO:BP terms among baseline-divergent genes mapped to brain-expansion metaterms, compared to 33.2% among divergent rGenes (Fig. 5D), localizing functionally relevant divergence to stimulus-dependent responses.

**Fig. 5.**
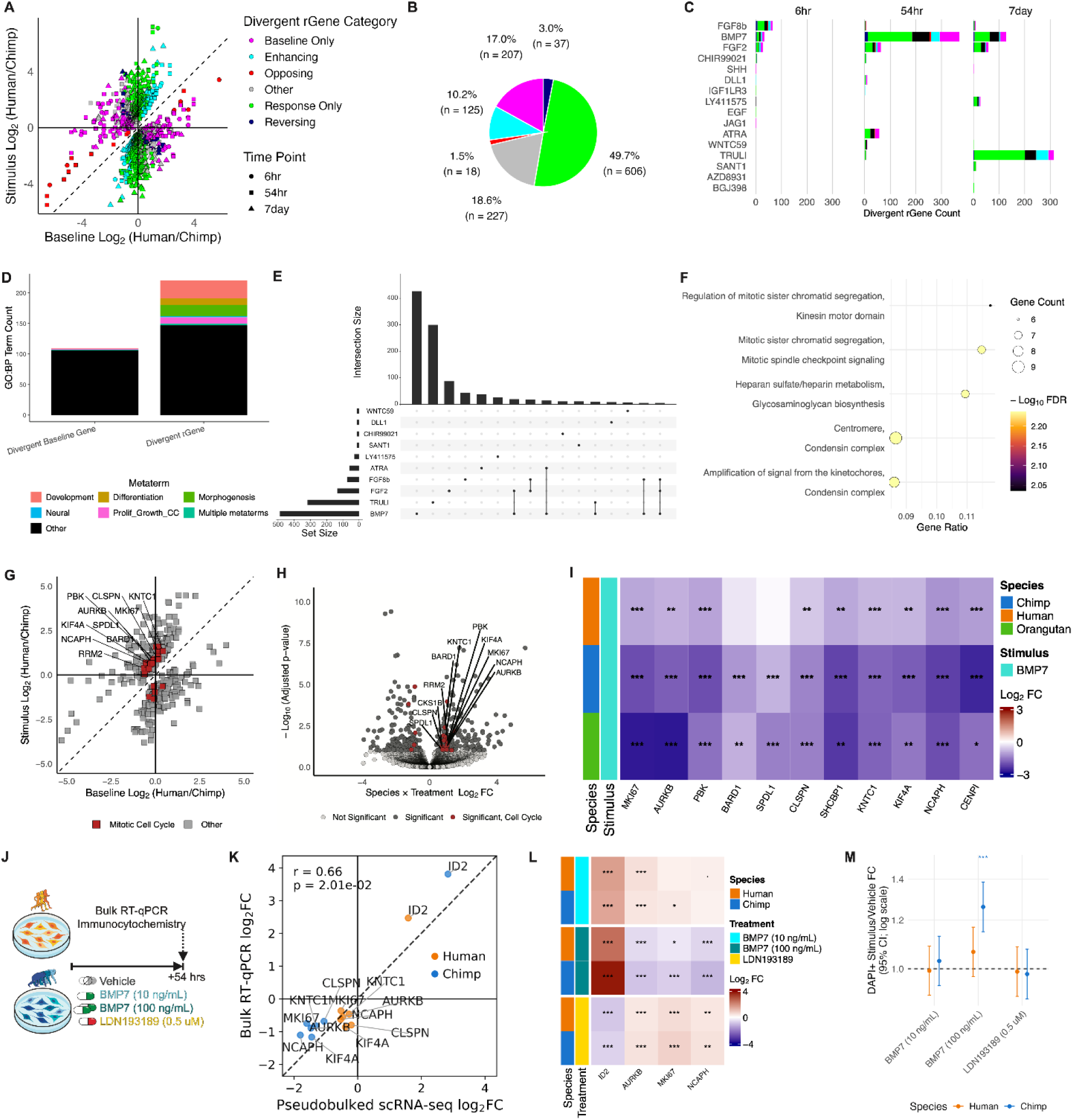
Stimulus-dependent regulatory divergence reshapes conserved signaling responses. (A–B) Classification of divergent response genes (rGenes) based on how stimulus exposure modifies species differences in gene expression. Divergence may emerge only under stimulation, be enhanced or opposed by stimulation, or be independent of stimulus context. **(C)** Distribution of divergent rGene categories across stimuli and time points, showing that divergence is concentrated in specific signaling conditions. **(D)** Functional enrichment comparison between baseline-divergent genes and stimulus-dependent divergent rGenes, revealing increased representation of developmental and proliferation-related processes among stimulus-dependent divergence. **(E)** Overlap of divergent rGenes across stimuli, indicating stimulus-specific and shared components of regulatory divergence. **(F)** Protein interaction enrichment among divergent BMP7 rGenes at 54 hrs, highlighting coordinated perturbation of interacting gene products. **(G–H)** Species divergence in BMP7 responses at 54 hrs, highlighting enrichment of mitotic cell cycle genes among divergent rGenes. Dots represent divergent rGenes (FDR < 0.10) colored by membership under the parent GO:BP term ‘Mitotic cell cycle’. **(I)** Cross-species comparison of BMP7 responses at 54 hrs across species for divergent mitotic cell cycle genes, showing a consistent pattern in which human cells exhibit a reduced decrease in expression relative to chimpanzee and orangutan. **(J)** Experimental design for validation of divergent BMP7 responses using bulk RT–qPCR and ICC. **(K)** Concordance between pseudobulk scRNA-seq and bulk RT–qPCR measurements of divergent BMP7 responses. Conserved BMP7 rGene *ID2* was excluded from the correlation calculation to restrict analysis to divergent rGenes. **(L)** Gene expression responses to BMP7 pathway modulation by bulk RT-qPCR across divergent rGenes associated with the mitotic cell cycle, with conserved BMP7 rGene *ID2* included as a positive control. **(M)** Quantification of cellular responses to BMP7 pathway modulation in intraspecies cultures, showing species differences in stimulus-dependent changes in cell number. Confidence intervals represent 95% intervals on log-transformed values.

We therefore focused on species divergence within the morphogen response contexts. Divergent rGene counts increased over time (139 genes at 6 hrs to 548 genes at 7 days) and correlated with conserved rGene counts but not cell number differences between species (Fig. S8D-G). Notably, 91.7% of divergent rGenes were detected in only one stimulus condition (Fig. 5E).

We next decomposed species divergence into baseline and stimulus-dependent components, and their interaction. We categorized divergent rGenes by how stimulus response affected baseline species divergence (Fig. 5A): genes divergent only in response (response only, 49.7%, 601 genes) or only at baseline (baseline only, 17.0%, 207 genes), and genes whose baseline divergence was amplified (enhancing, 10.2%, 125 genes), diminished (opposing, 1.5%, 18 genes) or reversed direction (reversing, 3%, 37 genes) by the stimulus (Fig. 5A-B,Table S9). More than half of species differences in rGenes were detectable only in the response context.

To dissect the functional relevance of divergent rGenes, we focused on FGF8b, FGF2, and BMP7, three stimuli with the strongest species divergence (Fig. 5C). At 6 hrs, FGF8b and FGF2 treatments showed the most divergent rGenes (66 and 28, respectively), with nine shared human-upregulated genes consistent with human-derived expression patterns (Fig. S8H-J). The most significant human-specific differences at 6 hrs included *PDE4B*, *GAP43*, *DNMBP*, *DST*, *DLGAP1*, and *ST6GALNAC5* (Fig. S8I). Species divergence in *PDE4B*, a phosphodiesterase that regulates cAMP levels via its degradation, increased more than 6-fold in response, while divergence in *DNMBP*, a guanine nucleotide exchange factor for CDC42, increased more than 2-fold in response and was detected only in the response context (Fig. S8H-I). At 54 hrs, *PDE4B*, *DST*, and *ST6GALNAC5* continued to show human-upregulated divergence. Several additional human-upregulated genes emerged at this timepoint, including *NPY*, *NRP1*, *IL17RD*, *LPAR1*, and *BMPR1A,* all associated with the MAPK and ERK1/2 signaling cascade, along with an enrichment for GPCR signaling (Fig. S8L-N). These observations are consistent with a proposed FGF-ERK-BMP signaling axis that expands mammalian cortical progenitor cells.^50^

We next focused on BMP7, which elicited the strongest and most coherent species divergence, to examine how early signaling differences propagate to downstream cellular and proliferative phenotypes. Among BMP7 divergent rGenes at 6 hrs human-upregulated genes were enriched for WNT signaling and proliferation, while chimpanzee-upregulated genes were enriched for cell junction assembly (Fig. S8O). Human-upregulated BMP7 rGenes included TFs mediating BMP and WNT responses (*SMAD3*, *SOX4*, and *GATA3)*, with species divergence in *SMAD3* and *GATA3* observable only in the response context (Fig. S8P-R). Consistent with increased BMP-WNT crosstalk in human telNECs, *NKD1*, a negative feedback inhibitor of canonical WNT signaling, responded more strongly to BMP7 in human than chimpanzee (Fig. S8P-R). Conversely, *ROR2*, a negative regulator of canonical WNT signaling, was upregulated in chimpanzee in response to BMP7, further amplifying existing baseline species divergence (Fig. S8P-R). Among chimpanzee-upregulated cell junction genes, *PARD6B,* a cortical hem marker involved in asymmetric cell division,^84^ showed species divergence in BMP7 response (Fig. S8R). Finally, *VEGFC*, associated with glial and epithelial cell proliferation, was human-upregulated in response to BMP7 and detected as a species difference only in the response context (Fig. S8R).

These early differences increased by 54 hrs, when human-upregulated BMP7 rGenes were far more numerous (Fig. 5C) and their protein products were enriched for mitotic cell cycle interactions (Fig. 5F). Cell cycle regulators *MKI67*, *AURKB*, and *KNTC1* were downregulated to a greater extent in chimpanzee and orangutan than in human, suggesting a derived human response to maintain their expression (Fig. 5G-I). For several of these genes, species divergence was observed only in response to BMP7. Human-upregulated BMP7 rGenes were also enriched for epithelial to mesenchymal transition (EMT) consistent with the EMT-like transition from NECs to radial glia, while chimpanzee-upregulated rGenes were enriched for morphogenesis and cytoskeletal terms associated with neuronal development (Fig. 8S-U). These results suggest that early divergence in signaling may propagate to downstream cellular phenotypes.

We validated these findings by bulk RT-qPCR on independent human and chimpanzee intraspecies cultures. We focused on eight divergent BMP7 rGenes associated with the cell cycle along with the conserved BMP7 rGene *ID2*. Following treatment with vehicle or BMP7 (100 ng/mL) (Fig. 5J), bulk and single-cell responses were well correlated (Pearson’s r = 0.66, *p* =2.01e-2, Fig. 5K). All cell-cycle divergent rGenes were downregulated across species in response to BMP7 with 5/6 decreasing less in human cultures, while *ID2* showed a robust species-conserved increase. *MKI67*, *AURKB*, and *NCAPH*, remain nominally significant (*p* < 0.10) for species divergence in response in the bulk assay (Fig. 5L).

Species divergence of *AURKB*, *MKI67*, and *NCAPH* required BMP signaling: these genes displayed a species-conserved increase in response to BMP receptor inhibitor LDN193189, and a low dose of BMP7 was not sufficient to unmask human-specific divergence (Fig. 5M). High-dose BMP7 increased culture density in human and chimpanzee intraspecies cultures, while low-dose BMP7 and LDN193189 did not (Fig. 5M). Combined with the species-conserved decrease in cell cycle gene expression (Fig. S7I), these observations suggest that BMP7-induced increases in cell density drive cell cycle downregulation, but that human telNECs are relatively resistant to this effect compared to chimpanzee and orangutan.

Together, these results show that, despite widespread baseline species differences, functionally relevant divergence emerges in a stimulus-dependent manner against a background of conserved signaling responses and propagates through downstream pathways to distinct cellular phenotypes.

### Evaluating the functional and evolutionary importance of rGenes

Genes regulated by developmental signaling during cortical expansion are predicted to be under strong evolutionary constraint. We therefore tested whether rGenes are enriched for gene sets reflecting evolutionary constraint and functional relevance relative to non-rGenes (two-tailed Fisher’s exact test throughout). Conserved rGenes were enriched for LoF-intolerant genes (OR = 1.63; FDR = 7.32e-34) and GWAS genes (OR = 1.58; FDR = 3.31e-37), and showed weaker enrichment for genes with genetically mapped cis-regulatory variation (eGenes) (OR = 1.12; FDR = 7.06e-4) (Fig. 6A). They were further enriched for protein-coding genes (OR = 2.15; FDR = 5.09e-70) and depleted for pseudogenes (OR = 0.43; FDR = 1.41e-10), antisense RNAs (OR = 0.65; FDR = 1.97e-04), and long non-coding RNAs (OR = 0.68; FDR = 1.03e-04) (Fig. 6B). Given the role of telNECs in cortical surface area expansion, we further examined links to neuroanatomical variation. Conserved rGenes were enriched for genes associated with cortical surface area (OR = 1.58; FDR = 0.015) and microcephaly (OR = 1.43; FDR = 8.9e-3) (Fig. 6C), consistent with the radial unit hypothesis.

**Fig. 6.**
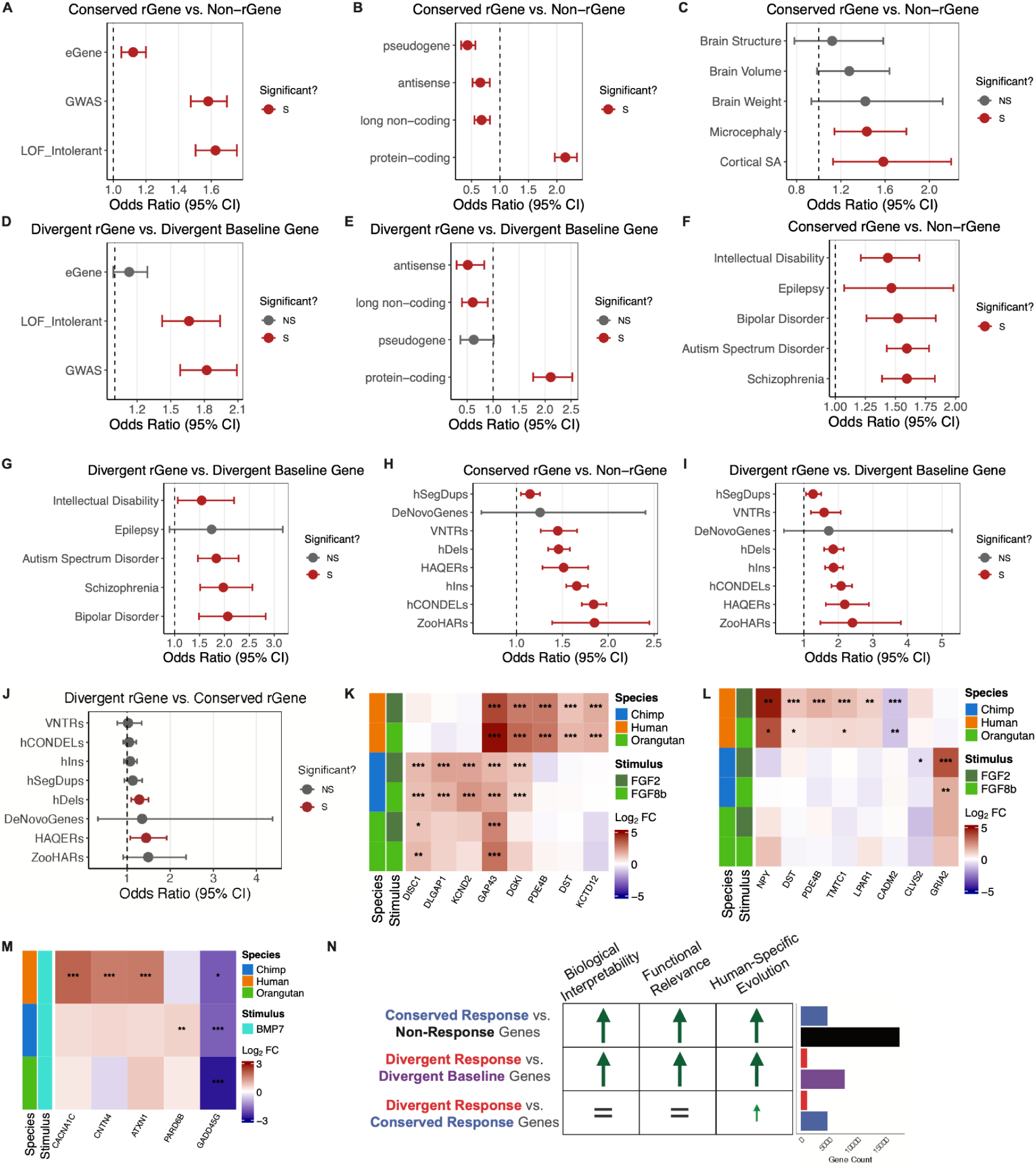
Stimulus-dependent responses are enriched for functionally constrained, disease-associated, and evolutionarily modified genes. (A–J) Forest plots show odds ratios (ORs) with 95% confidence intervals derived from two-sided Fisher’s exact tests. The dashed vertical line indicates no enrichment (OR = 1). Nominal enrichments (p ≤ 0.05) are shown in red; non-significant results are shown in gray. **(A–C)** Conserved rGenes are enriched for LoF-intolerant, protein-coding, and GWAS-associated genes (A), protein-coding genes and depleted for non-coding categories (B), and genes associated with neuroanatomical traits relevant to cortical expansion, including microcephaly and cortical surface area (C) relative to non-rGenes. **(D–E)** Divergent rGenes are enriched for functionally constrained (D) and protein-coding genes and depleted for non-coding categories (E) relative to baseline-divergent genes. **(F–G)** Both conserved (F) and divergent (G) rGenes are enriched for NDD-associated genes, including ASD, schizophrenia, bipolar disorder, intellectual disability, and epilepsy. **(H–J)** Enrichment of genes overlapping or proximal to human-specific genomic features. rGenes are enriched across multiple feature classes, including conserved rGenes relative to non-rGenes (H), with divergent rGenes showing preferential enrichment relative to conserved rGenes near specific feature classes (J). **(K–M)** Examples of divergent rGenes linking stimulus-dependent regulatory divergence to human-specific sequence variation and neuropsychiatric disease. Heatmaps show cross-species gene expression responses of telNECs to FGF8b and FGF2 at 6 hrs (K) and 54 hrs (L), and BMP7 at 6 hrs (M), for divergent rGenes overlapping or proximal to human-specific genomic features (hDels or HAQERs) and associated with ASD, schizophrenia, or bipolar disorder. **(N)** Summary of enrichment patterns across gene categories. Stimulus-dependent responses (both conserved and divergent) show increased biological interpretability, functional relevance, and enrichment for human-specific evolutionary features relative to baseline comparisons, with divergent responses showing the strongest enrichment for human-specific sequence variation.

We next asked whether stimulus-dependent divergence similarly captures functionally meaningful regulatory variation. Divergent rGenes were enriched for signatures of functional importance relative to baseline-divergent genes. These included LoF-intolerant genes (OR = 1.66; FDR = 1.92e-10), protein-coding genes (OR = 2.11; FDR = 1.97e-18), and GWAS genes (OR = 1.82; FDR = 1.02e-16), but not eGenes, and showed depletion of antisense (OR = 0.51, FDR = 1.28e-02) and long non-coding RNAs (OR = 0.61, FDR = 1.79e-02; two-tailed FET) (Fig. 6D–E).

Given proposed tradeoffs between human brain evolution and NDD risk,^92^ we tested whether rGenes were enriched for NDD-associated genes. Both conserved rGenes (versus non-rGenes) and divergent rGenes (versus baseline-divergent genes) were enriched for ASD, SCZ, BPD, intellectual disability, and epilepsy-associated genes (Fig. 6F–G). Direct comparison of divergent and conserved rGenes revealed no additional enrichment for ASD or SCZ, with only BPD, a disorder linked to prenatal brain development and developmental signaling pathways,^93^ showing nominal enrichment among divergent rGenes (OR = 1.47, p = 0.021), driven by responses to BMP7 and FGF2 (Table S10). These results indicate that developmental signaling pathways dynamically regulate NDD-linked genes in telNECs, with both conserved and species-divergent responses contributing to this regulation.

To connect rGenes to sequence-level evolution, we tested enrichment near human-specific genomic features, including accelerated regions, structural variants, and repetitive elements (ZooHARs,^94^ HAQERs,^95^ hCONDELs,^96^ hDels,^97^ hIns,^97^ hSegDups, VNTRs^98^ and *de novo* genes^99^). With the exception of *de novo* genes, for which few features exist, both conserved and divergent rGenes were significantly enriched across these feature classes relative to non-rGenes and baseline-divergent genes, respectively (Fig. 6I-J). Divergent rGenes showed additional enrichment for hDels and HAQERs compared to conserved rGenes (Fig. 6J). In total, 639 divergent rGenes were proximal to human-specific genetic features, including 161 (25.2%) linked to NDD risk, connecting stimulus-dependent regulatory divergence to human-specific sequence evolution and disease.

A subset of divergent FGF rGenes influence cAMP–PKA signaling and overlap with human-specific sequence variation and disease-relevant pathways. In response to FGF, Disrupted in Schizophrenia 1 (*DISC1*) showed a human-specific reduction by 6 hrs compared to chimpanzee and orangutan telNECs (Fig. 6K). DISC1 interacts with PDE4B, another early divergent FGF rGene and an ASD/SCZ risk gene, to regulate intracellular cAMP levels via PKA signaling.^100^ *PDE4B* itself overlaps a human-specific segmental duplication and lies near additional human-specific variants. At 54 hrs, Neuropeptide Y (*NPY*), a SCZ-linked gene that reduces cAMP-PKA signaling through GPCR-coupled inhibition of adenylyl cyclase,^101^ showed human-specific upregulation, further implicating this pathway. Together, these observations implicate cAMP–PKA signaling as a candidate mediator of species differences in cortical progenitor regulation, consistent with prior work linking ERK–cAMP–PKA signaling to cortical expansion.^50^

Among early divergent BMP7 rGenes, *CACNA1C* provides a complementary example linking stimulus-dependent regulatory divergence to human-specific sequence changes with functional consequences. BMP7 enhanced baseline expression divergence of *CACNA1C*, a BPD and SCZ risk gene encoding the pore-forming subunit of the Ca_V_1.2 calcium channel. *CACNA1C* harbors an intronic human-specific tandem repeat expansion TRACT^98,102–104^ that acts as an enhancer in neural progenitors to increase *CACNA1C* expression,^102^ and modulates activity-dependent responses in neurons, including by repression of the cell cycle regulator *GADD45G.*^104^ Human telNECs respond to BMP7 with increased *CACNA1C* expression across time points, in addition to increased baseline expression at 6 hrs and 54 hrs (Fig. 6m). These examples suggest that human-specific regulatory changes may reshape morphogen responses across signaling contexts, shifting downstream signaling activity that regulates progenitor state and proliferation.

Collectively, these results show that stimulus-dependent divergence preferentially targets functionally constrained, disease-associated, and evolutionarily modified genes, linking developmental signaling to human brain expansion and neuropsychiatric risk.

## DISCUSSION

The expansion of the human brain likely arose from genetic changes that altered developmental programs in telencephalic progenitors. Because species differences in brain size are evident by mid-gestation,^1^ these changes likely originate at the neuroepithelial stage and compound across development.^2,3^ Yet identifying the relevant regulatory changes has been challenging because baseline gene regulatory divergence is shaped by neutral drift and shows limited overlap with variants influencing complex traits.^13,17,18^ We therefore asked whether functionally relevant regulatory divergence is preferentially revealed in response to developmental signaling. By profiling transcriptomic responses of telNECs to morphogens, we sought to identify the “missing regulation” linking sequence divergence to progenitor behavior relevant to brain expansion.

Across 20 perturbations spanning nine signaling pathways, telNECs from human, chimpanzee, and orangutan showed broadly conserved transcriptional and cellular responses, with modulation of FGF, BMP, and Notch signaling producing the strongest effects. Conserved response genes were enriched for processes related to cortical expansion, including genes linked to microcephaly, cortical surface area variation, and NDDs, emphasizing the functional importance of morphogen-responsive genes. Against this conserved background, species-divergent responses were frequently detectable only under stimulation: more than half of divergent rGenes were absent from baseline comparisons. Despite representing seven-fold fewer genes than baseline divergent genes, these stimulus-dependent differences showed stronger enrichment for developmental functions relevant to brain expansion, including resistance to BMP7-induced high culture density-associated cell cycle downregulation. Thus, baseline divergence is widespread but weakly informative, whereas stimulus-dependent divergence is sparse, structured, and functionally prioritized.

Consistent with this prioritization, both conserved and divergent rGenes were enriched for NDD-associated genes and for proximity to human-specific genomic features, with divergent rGenes showing stronger enrichment near hDels and HAQERs. These enrichments suggest that stimulus-dependent regulatory divergence preferentially targets genes under evolutionary constraint with roles in human brain development.

Several convergent observations support a model in which human telNECs have not acquired new signaling pathways, but instead exhibit altered thresholds within conserved signaling networks. Against a background of broadly conserved responses, human telNECs showed relative resistance to BMP7-driven cell cycle downregulation, human-specific changes in FGF-dependent cAMP–PKA signaling, and BMP7-induced cross-activation of WNT targets. These pathways jointly regulate progenitor self-renewal, developmental progression, and differentiation timing.^105^ Although these regulatory relationships are deeply conserved, fate transitions are protracted in humans. This framework is consistent with prolonged maintenance of human progenitors under sustained growth factor support,^60^ human gains of enhancers near growth-factor and cell-cycle genes,^106^ and increased robustness of human progenitors to perturbation of G1/S regulators.^107^ In this view, stimuli reveal divergence because baseline conditions do not sufficiently challenge these thresholds. When morphogen input is varied, human cells resist transitions that more readily advance chimpanzee and orangutan progenitors.

Gene-level examples illustrate how stimulus-dependent divergence manifests at key nodes within conserved signaling networks. In response to FGF, *DISC1* showed a human-specific reduction, while *PDE4B*—an interacting ASD/SCZ risk gene overlapping a human-specific segmental duplication—and *NPY*, a regulator of cAMP signaling, showed human-specific divergence, collectively implicating altered cAMP–PKA signaling. In the BMP7 response, *CACNA1C*, a BPD/SCZ risk gene harboring a human-specific enhancer-like tandem repeat expansion, showed elevated baseline expression and further induction in human cells, consistent with altered activity-dependent regulation and downstream effects on cell cycle genes such as *GADD45G*. Together, these examples suggest that human-specific regulatory changes may shift how conserved signaling inputs are interpreted.

These findings demonstrate that systematic delivery of physiological stimuli unmasks functionally relevant gene expression divergence that is invisible in steady-state comparisons.^22,23^ The overlap of rGenes with signatures of function, selection, and disease in telNECs extends stimulus-response approaches applied in other cell types, contexts, and timescales^24–26,28–36,108–114^ to human brain evolution and neuropsychiatric vulnerability, providing a framework for dissecting evolved developmental signaling responses across the hominid lineage.

This study has several limitations. We examined acute perturbations at a single dose across a defined set of pathways; additional pathways, doses, durations, and combinations may reveal further divergence. Our analyses focused on telNECs, and later progenitor states will be required to determine how early divergence propagates through cortical development. Finally, single-cell transcriptomics does not capture post-transcriptional or post-translational regulation,^115^ and the relative contributions of cis- and trans-regulatory changes remain unresolved, requiring multi-omic and comparative functional approaches.^116–119^ These findings suggest that functionally relevant regulatory divergence linking genetic variation to human brain expansion is preferentially revealed through context-dependent responses to developmental signaling.

---

**Fig. S1.**
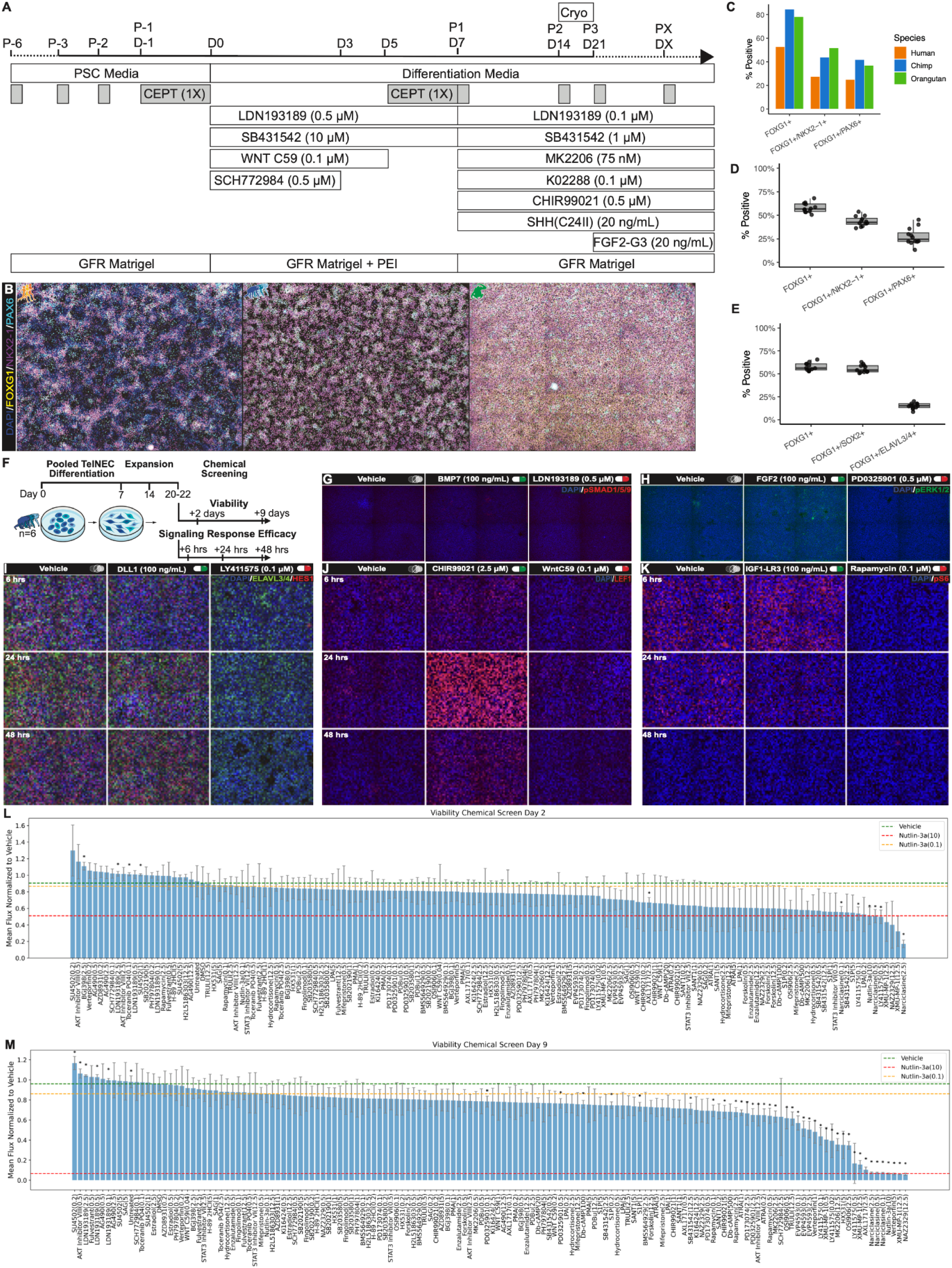
Directed differentiation of telNECs and validation of chemical screening framework. **(A)** Schematic of PSC maintenance, telNEC directed differentiation, and expansion protocol. Cryo, cryopreservation; GFR, growth factor reduced. **(B–C)** Immunocytochemistry (ICC) and quantification at day 43 demonstrating robust generation of telencephalic progenitors across multi-individual, interspecies cultures of human and chimpanzee and single-individual culture of orangutan. **(D–E)** Quantification of ICC from interspecies pooled cultures at day 54 (+6 hrs), corresponding to Fig. 1E–F, respectively, confirming reproducible regional and neurogenesis marker expression across biological replicates. Four technical replicate images were collected from each of three biological replicate wells. **(F)** Experimental design of chemical screening framework, including signaling response assays (G–K and Fig. 1H–I) and viability time-course assays (L–M) . **(G–K)** ICC-based validation of pathway activation following chemical perturbation. Panels show activation of BMP (pSMAD1/5/9) (G), FGF (pERK1/2) (H), NOTCH (HES1; with ELAVL3/4 marking neuronal differentiation) (I), WNT (LEF1) (J), and mTOR (pS6) (K) signaling pathways across time points. **(L–M)** Chemical viability screen across candidate compounds and doses at 2 days (L) and 9 days (M), showing differential effects on culture viability. Bars represent mean flux normalized to vehicle; error bars indicate standard deviation. Asterisks denote FDR < 0.10. Nutlin-3a is included as a positive control for p53 activation.

**Fig. S2.**
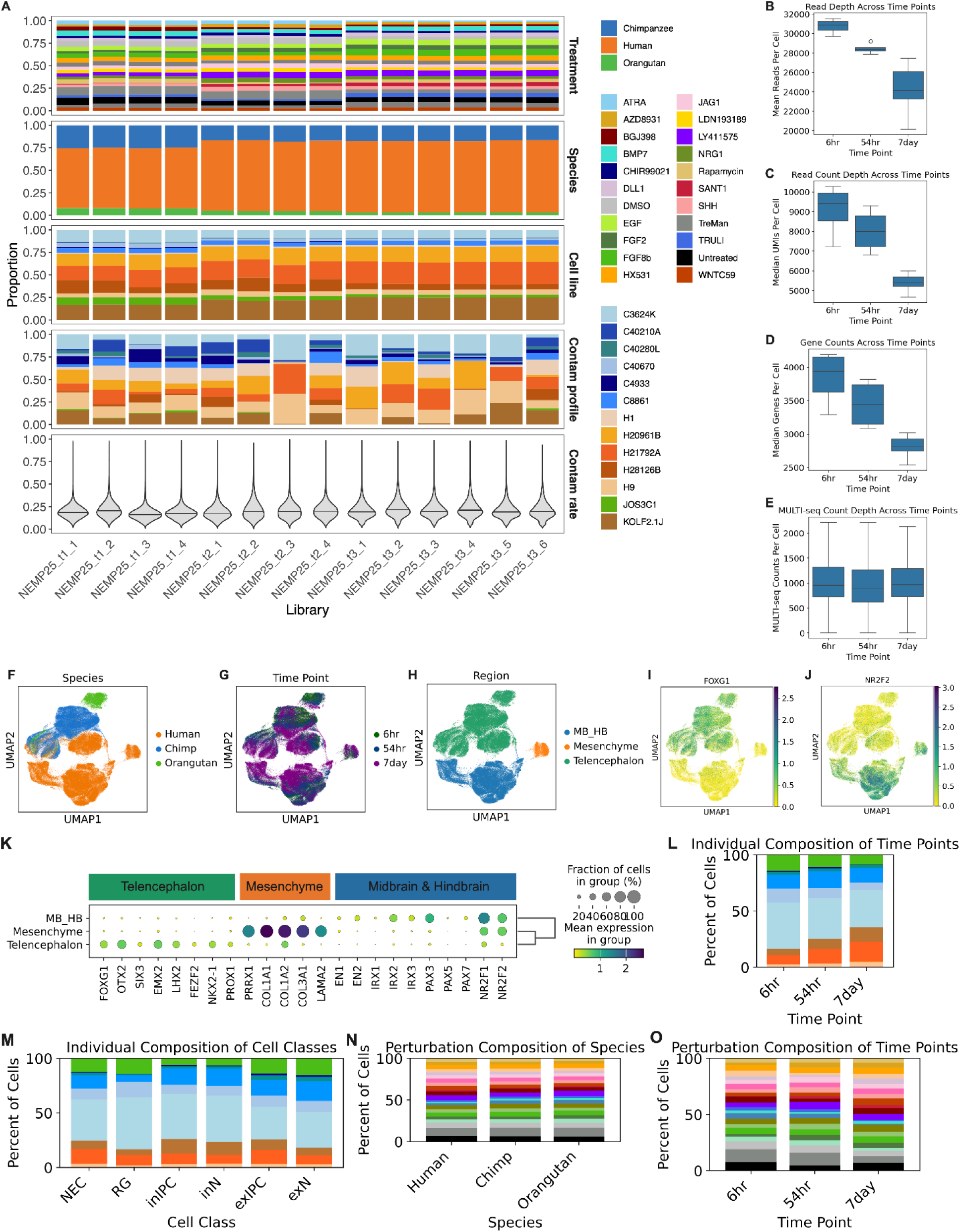
Demultiplexing accuracy, sequencing quality, and validation of telencephalic identity across pooled single-cell datasets. **(A)** Composition of sequencing libraries across perturbation, species, individual (cell line), and inferred ambient RNA contamination profiles, along with estimated contamination rates, demonstrating successful demultiplexing of pooled samples across conditions and time points. Library suffixes t1, t2, and t3 denote 6 hr, 54 hr, and 7 day time points, respectively. **(B–E)** Sequencing quality metrics across time points, including mean read depth (B), median read count per cell (C), median genes detected per cell (D), and MULTI-seq barcode read depth (E), indicating consistent data quality across libraries and experimental conditions. **(F–J)** UMAP embeddings of all cells prior to dataset integration, colored by species (F), time point (G), annotated region (H), expression of regional markers, including the telencephalon marker *FOXG1* (I) and off-target marker *NR2F2* (J). **(K)** Dot plot of regional marker expression across annotated regions, supporting anatomical annotation of cell populations. **(L–O)** Composition of cells across individuals, cell classes, perturbations, and time points, demonstrating balanced representation across key experimental variables and minimizing confounding in downstream analyses.

**Fig. S3.**
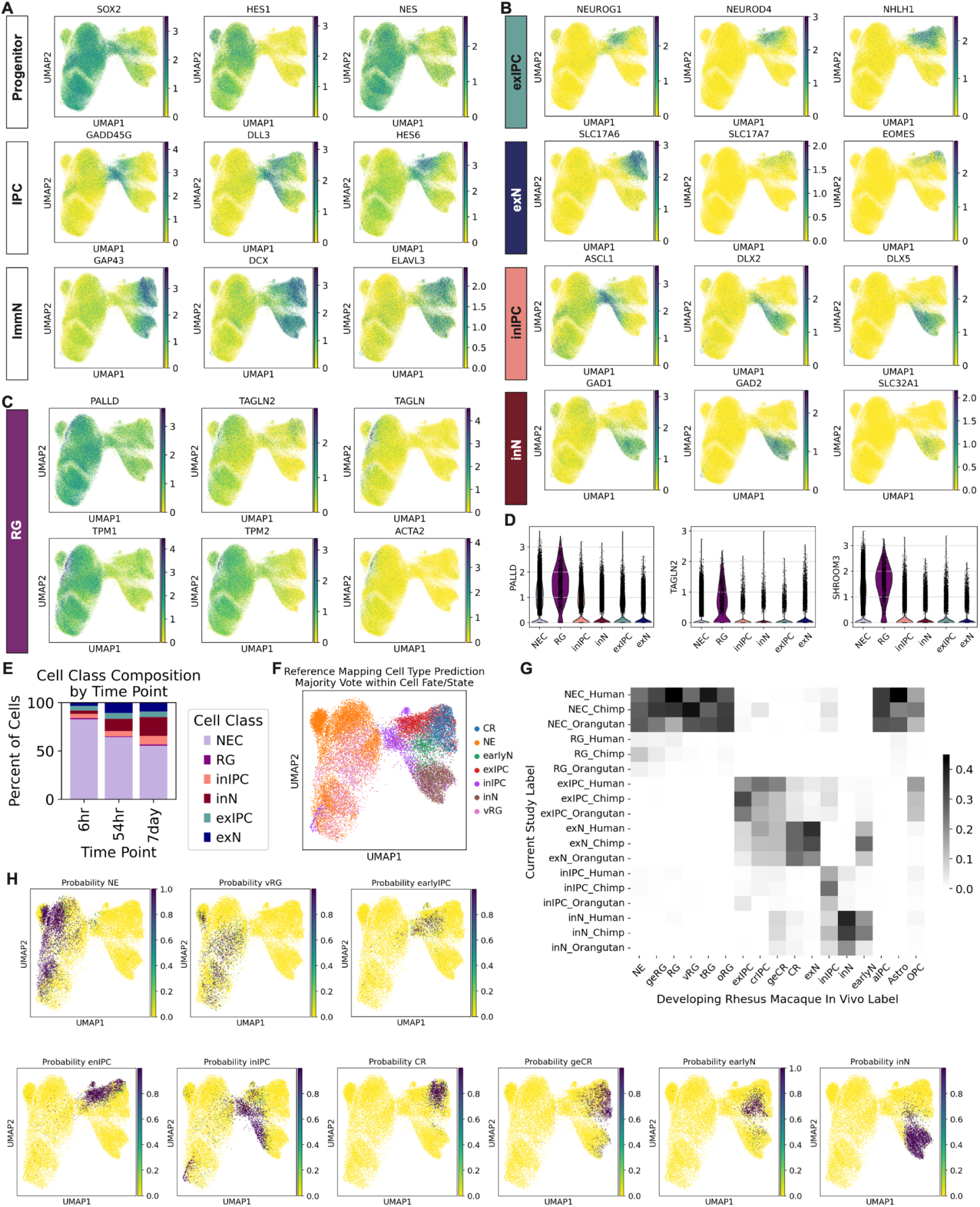
Cell class annotations and reference atlas cell type predictions. (A–B) UMAP plots showing expression of canonical marker genes defining major progenitor (NEC, RG), intermediate progenitor (IPC), and neuronal (immN) classes (A), as well as excitatory and inhibitory lineage markers (B), supporting cell class annotation across developmental trajectories. **(C)** UMAP plots of ventricular radial glia (vRG) marker genes (*PALLD*, *TAGLN2*) and cytoskeleton-associated genes, identifying a vRG-like population and transitional states between NEC and RG. **(D)** Expression of vRG markers (*PALLD*, *TAGLN2*) and NEC-to-RG transition marker *SHROOM3* across cell classes. **(E)** Cell class composition across time points, indicating expected progression from progenitor-dominated to more differentiated states. **(F)** UMAP colored by majority-vote reference atlas cell type predictions within annotated clusters, showing alignment between in vitro cell states and in vivo reference identities. **(G)** Confusion matrix comparing annotated cell classes with reference atlas cell type predictions across species, demonstrating concordant mapping between datasets. **(H)** UMAP plots of reference atlas prediction probabilities, illustrating spatial localization and confidence of cell type assignments across the manifold.

**Fig. S4.**
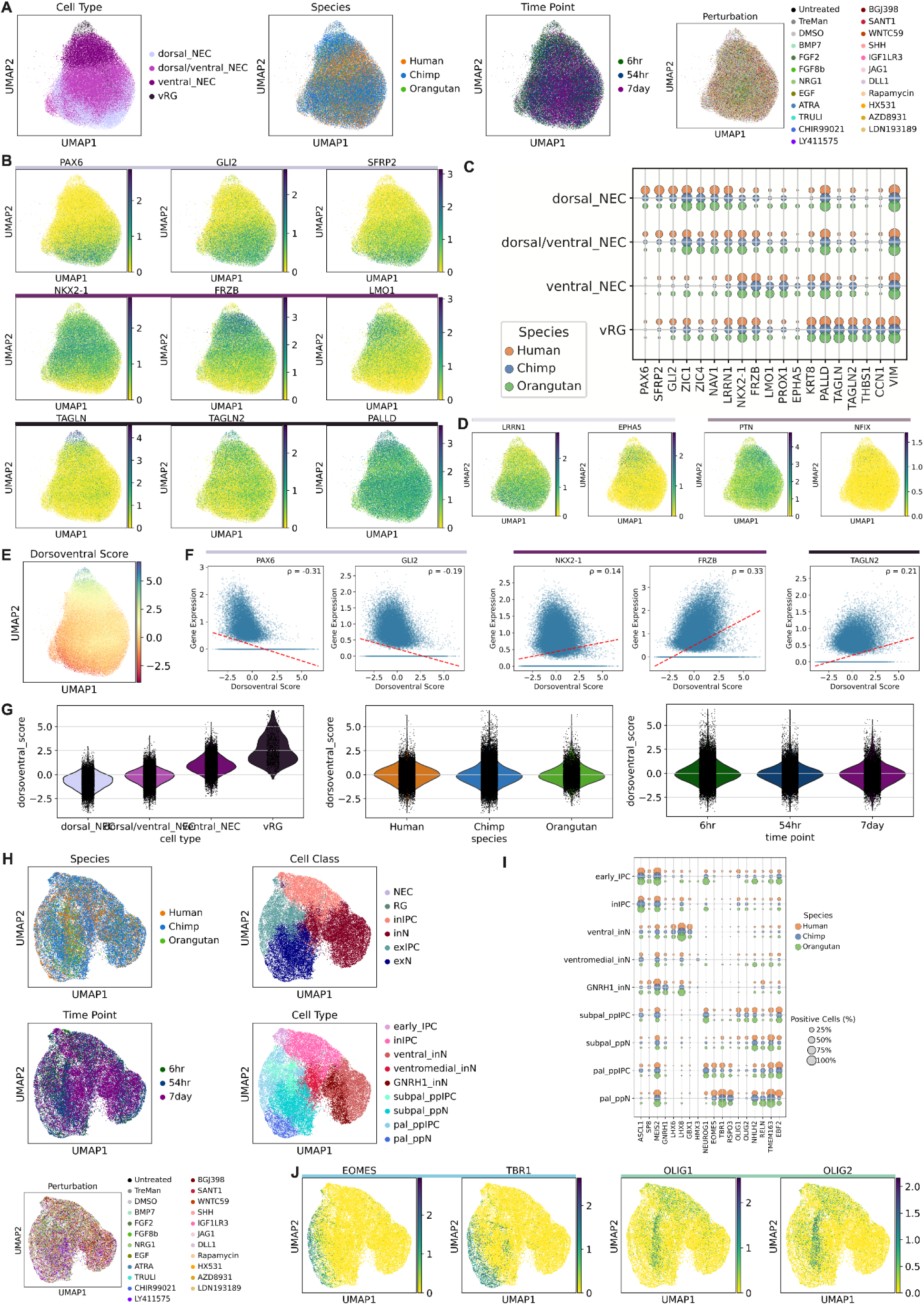
Annotation of progenitor and neuronal subtypes across telencephalic development. (A–C) UMAP of progenitor cells showing annotation of dorsal NEC, ventral NEC, and ventricular radial glia (vRG) states (A), expression of canonical marker genes (e.g., *PAX6, GLI2, SFRP2, NKX2-1, TAGLN2, PALLD*) (B), and distribution of marker expression across species (C), supporting identification of progenitor subtypes across conditions. **(D–F)** Analysis of the NEC-to-radial glia (RG) transition. Expression of transition-associated genes (*LRRN1, EPHA5, PTN, NFIX*) across UMAP space (D) and dorsoventral score gradients (E) reveal a continuum of progenitor regionalization after controlling for cell cycle effects. Correlation of gene expression with dorsoventral score (F) shows a shift from dorsal NEC markers (e.g., *PAX6*) to ventral NEC markers (e.g.*NKX2-1*) as well as upregulation of vRG-associated genes (e.g., *TAGLN2*). A small cluster transcriptionally similar to NECs but enriched for early vRG markers (*PALLD*, *TAGLN2*) and cytoskeleton-related genes, including SHROOM3, is observed, consistent with cells undergoing the NEC-to-vRG transition. **(G)** Distribution of dorsoventral scores across cell type, species, and time point, indicating consistent regionalization along the dorsoventral axis across experimental conditions. **(H–I)** UMAP and marker analysis of IPC and immature neuronal populations, including cell class annotations (H) and marker gene expression across subtypes (I), supporting classification of excitatory and inhibitory lineages. **(J)** UMAPs showing expression of dorsal (*EOMES, TBR1*) and ventral (*OLIG1, OLIG2*) neuronal markers, confirming appropriate regional patterning of neuronal subtypes. Together, these analyses support a continuous dorsoventral axis with comparable regional identity across species and time points.

**Fig. S5.**
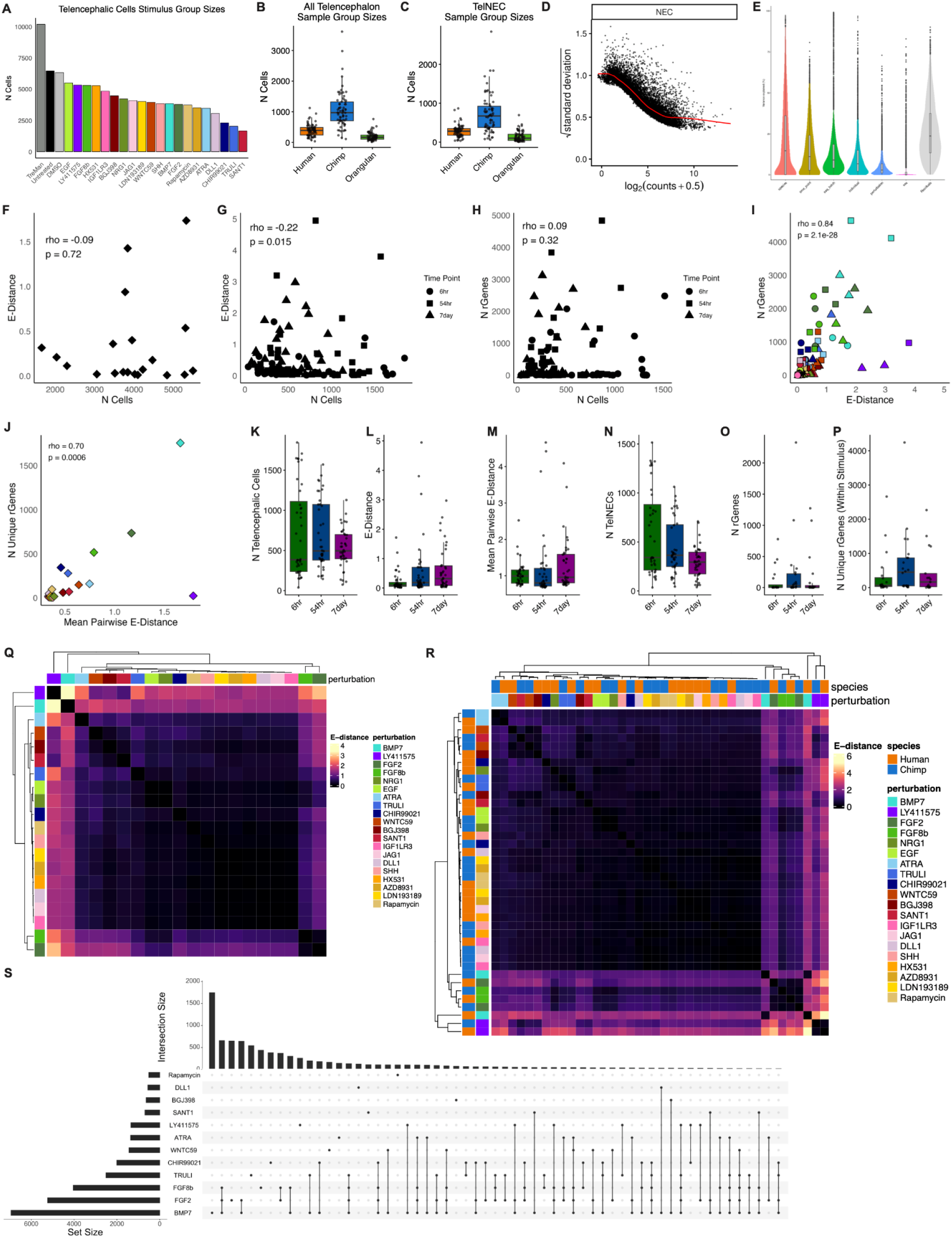
Robustness of morphogen response effect sizes and their relationship to rGene detection. (A–C) Distribution of stimulus and sample group sizes used for E-distance and differential expression analyses across telencephalic cells (A–B) and telNECs (C), establishing adequate sampling across conditions, species, and time points. **(D–E)** Model diagnostics for pseudobulk scRNA-seq analysis, including mean–variance relationships (voom; D) and variance partitioning across experimental variables (E), supporting appropriate modeling of gene expression variability. **(F–H)** Relationship between sample size and response metrics. E-distance shows minimal dependence on stimulus or sample group size (F–G), and the number of detected rGenes is not strongly driven by sample size (H), with most conditions yielding detectable rGene sets (≥10 genes in 112 of 124 conditions), indicating that response magnitude and rGene detection are not artifacts of sampling depth. **(I–J)** Relationship between global transcriptional response magnitude and gene-level divergence. E-distance is positively associated with the number of rGenes across sample groups (I) and with the number of unique rGenes across stimuli (J), linking overall response strength to the extent of differential gene regulation. **(K–P)** Distribution of sample sizes, E-distances, and rGene counts across time points, showing consistent response magnitudes and gene-level effects across 6 hr, 54 hr, and 7 day conditions. **(Q–R)** Pairwise E-distance heatmaps across stimulus groups (Q) and across species within perturbation groups (R), revealing structured relationships among morphogen responses and consistency across species. **(S)** Upset plot of rGene overlap across stimuli, showing both shared and stimulus-specific gene response programs.

**Fig. S6.**
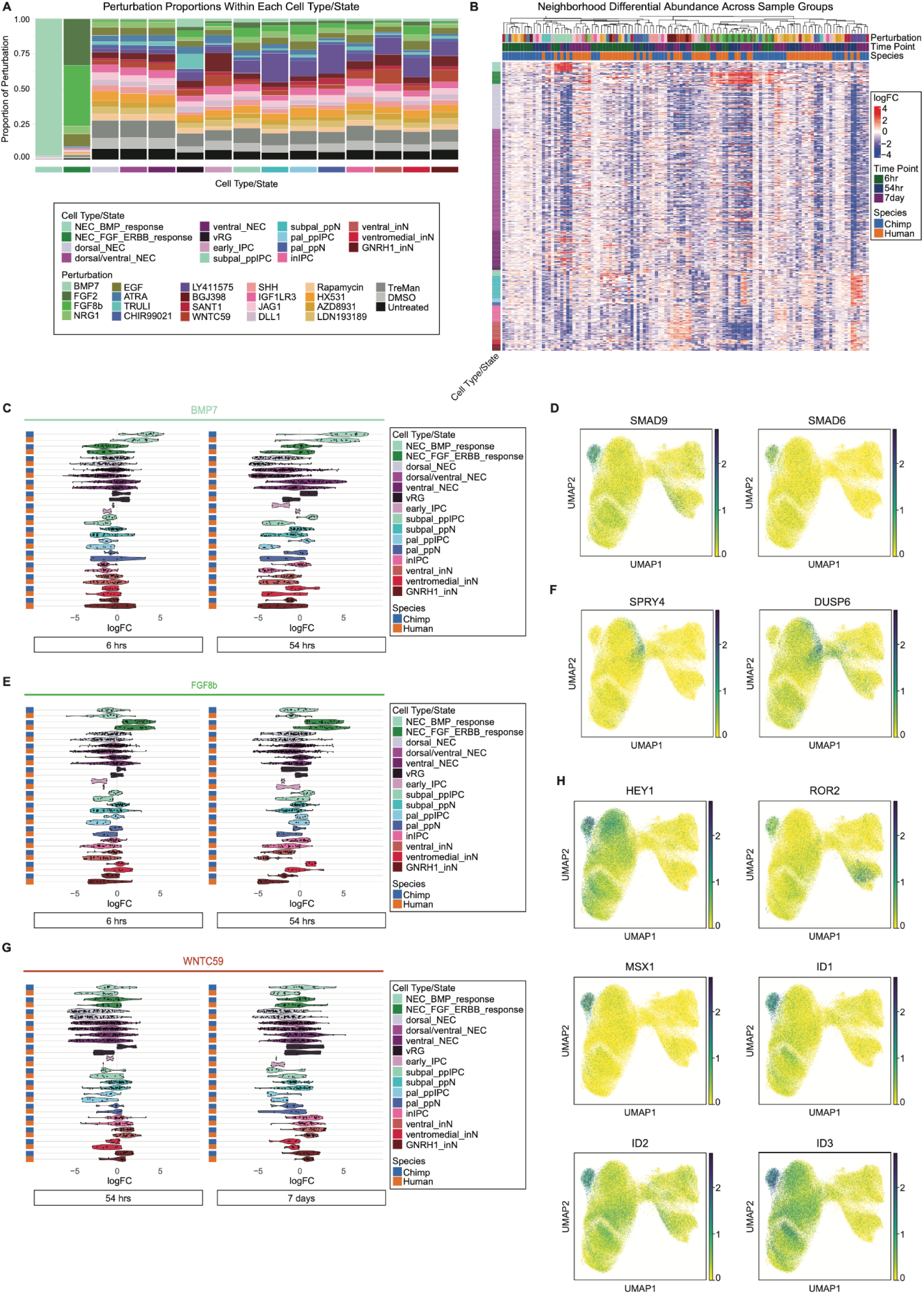
Cellular abundance responses to morphogens are structured across cell types and largely conserved across species. (A–B) Global structure of cellular abundance responses. Perturbations are broadly distributed across cell types and states (A), and neighborhood-level differential abundance changes show structured variation across sample groups (B), indicating coordinated shifts in cellular composition. **(C)** Cellular abundance responses to BMP7 across cell types and time points. Violin and strip plots show log fold changes in neighborhood abundance, with largely concordant responses between human and chimpanzee. **(D)** UMAP plots of canonical BMP pathway effectors (e.g., *SMAD9*, *SMAD6*), showing spatial patterns of pathway activation across cell states. **(E)** Cellular abundance responses to FGF8b across cell types and time points, demonstrating structured and largely conserved responses across species. **(F)** UMAP plots of canonical FGF/ERBB pathway effectors (e.g., SPRY4, DUSP6), highlighting pathway-aligned expression patterns. **(G)** Cellular abundance responses to WNTC59 across cell types and time points, showing cell-type-specific shifts that are broadly consistent across species. **(H)** UMAP plots of genes marking pallial and cortical hem identity in the Allen Institute Mouse Developing Brain ISH Atlas at E11.5, providing anatomical context for observed cellular abundance changes.

**Fig. S7.**
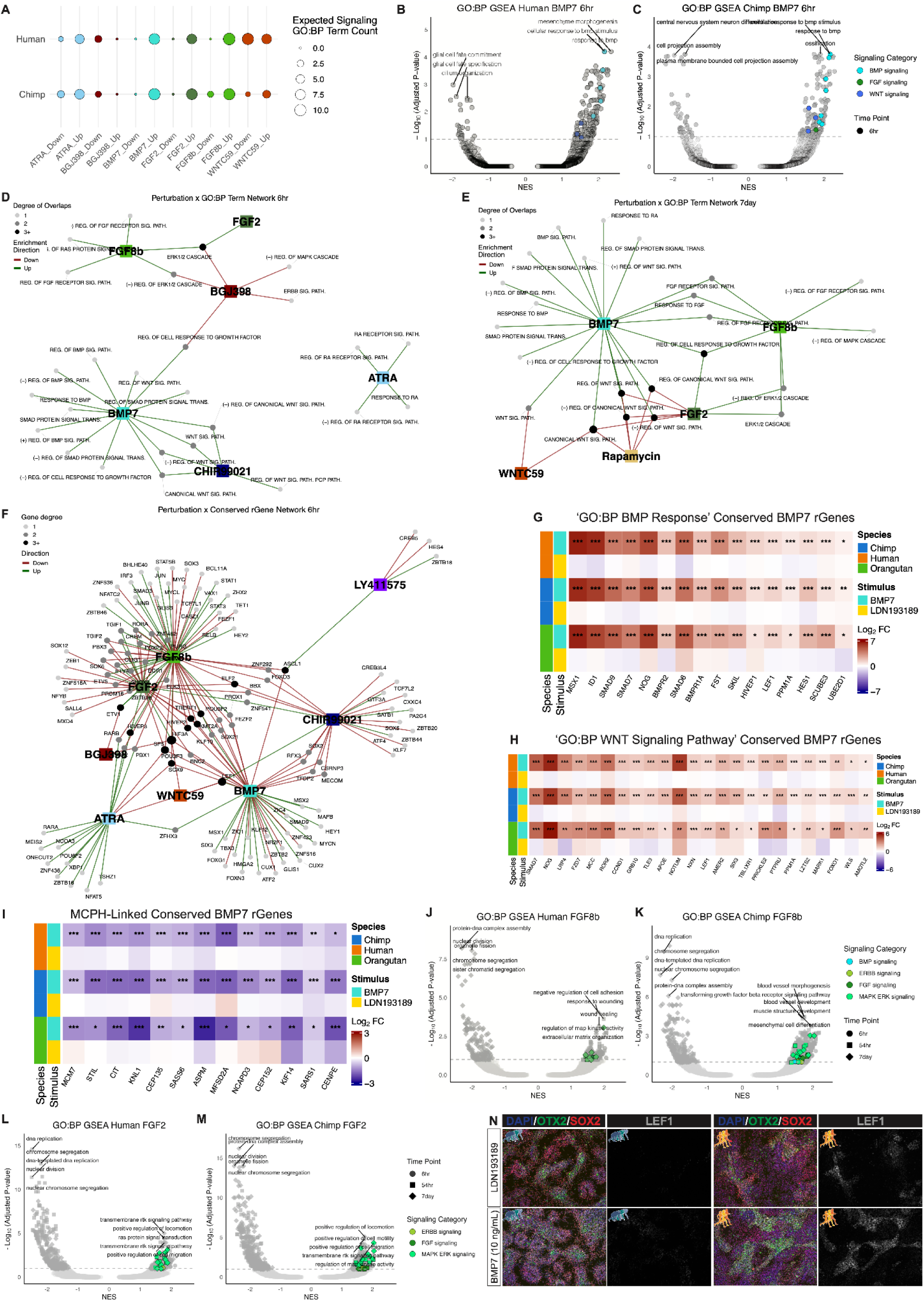
Specificity, dynamics, and mechanisms of conserved gene expression responses to morphogens. (A–C) Gene set enrichment analysis of conserved rGenes across species. Enriched GO:BP terms correspond to the perturbed signaling pathways, with consistent pathway-specific responses observed across species (A) and exemplified for BMP7 responses at 6 hrs in human (B) and chimpanzee (C). **(D–E)** Network of pathway enrichment across stimuli at early (6 hrs; D) and late (7 day; E) time points, with edges colored by direction of regulation. Enriched terms group by perturbed signaling pathway and show coordinated patterns of up- and down-regulation, with network structure evolving between early (6 hr) and late (7 day) time points. **(F)** Network of conserved rGene transcription factors annotated to signaling-related GO:BP terms, highlighting shared regulatory modules across perturbations. **(G–I)** Cross-species gene expression responses for conserved BMP7 rGenes at 54 hrs. Heatmaps show coordinated regulation of genes associated with BMP response (G), WNT signaling (H), and microcephaly-associated genes (I) by pseudobulk scRNA-seq modeling within telNECs, supporting pathway-specific and biologically relevant transcriptional programs, including cross-pathway activation of WNT-associated genes in response to BMP signaling. **(J–M)** Gene set enrichment analysis of conserved FGF8b (J) and FGF2 (K) responses for human (left) and chimpanzee (right) across time points, demonstrating consistent activation of FGF/MAPK-associated transcriptional programs. **(N)** ICC validation of conserved BMP7-responsive gene LEF1 across species under BMP activation (10 ng/mL BMP7)and inhibition conditions (0.5 µM LDN193189), confirming pathway-aligned transcriptional responses at the protein level.

**Fig. S8:**
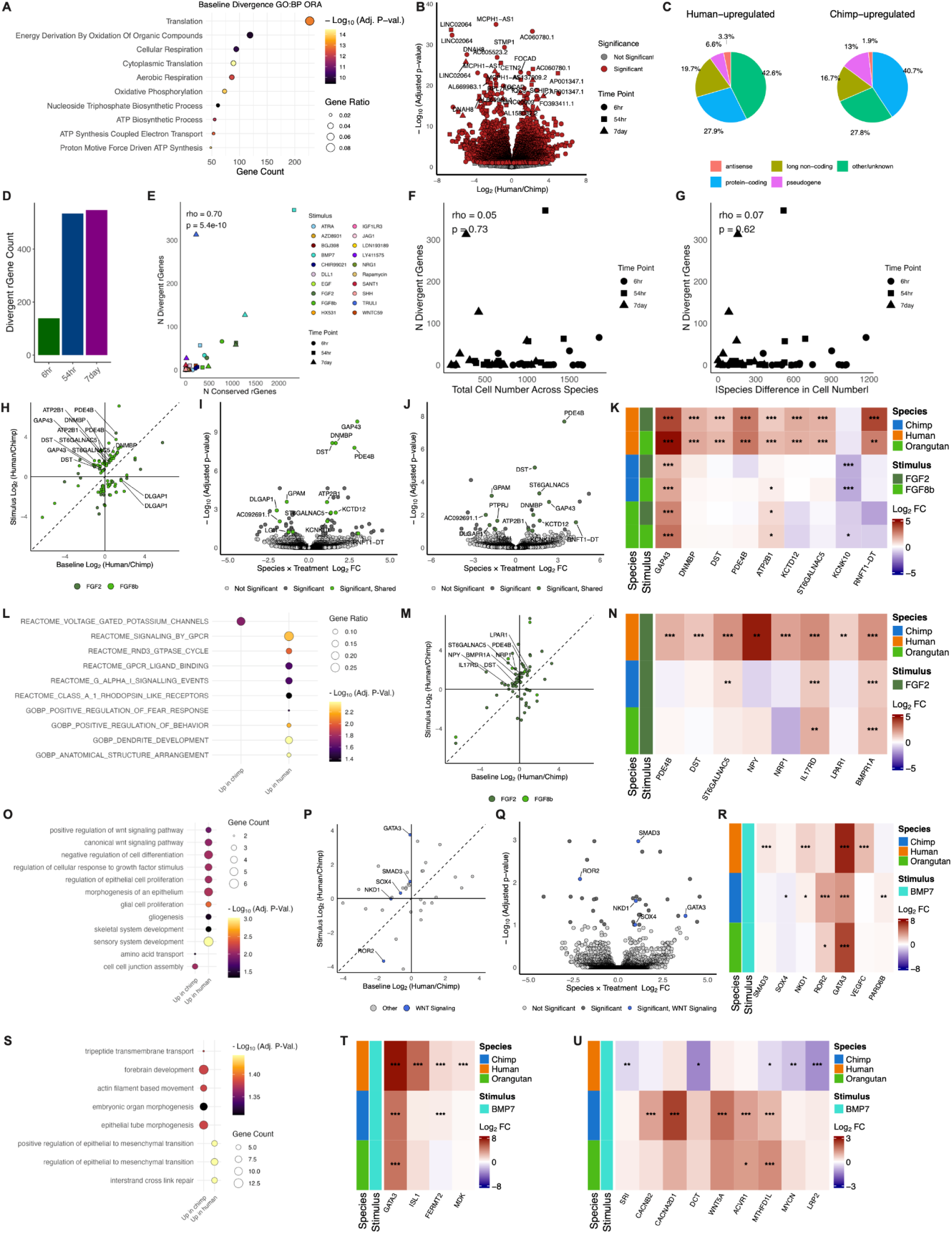
Baseline divergence is diffuse, whereas morphogen responses concentrate species differences into pathway-specific programs. (A–C) Baseline human–chimpanzee divergence is enriched for broad metabolic and cellular processes (A), with widespread but modest gene-level differences (B) spanning diverse gene biotypes (C). **(D–G)** Divergent rGene detection across sample groups. Divergent rGene counts increase over time (D) and scale with the number of conserved rGenes (E), but show minimal dependence on total cell number (F) or interspecies differences in cell number (G). **(H–K)** FGF responses at 6 hrs reshape baseline divergence. Divergent rGenes shift relative to baseline divergence (H) and show overlapping species-by-stimulus interaction effects across FGF8b (I) and FGF2 (J), with shared divergent responses highlighted in heatmaps (K) by pseudobulk scRNA-seq modeling. **(L–N)** At 54 hrs, FGF responses show pathway-specific enrichment. Divergent rGenes are enriched for signaling-related processes (L), exhibit consistent shifts relative to baseline divergence (M), and include genes associated with MAPK/ERK signaling (N). **(O–R)** BMP7 responses at 6 hrs reveal pathway-linked divergence. Divergent rGenes are enriched for developmental signaling processes (O), including canonical WNT signaling genes (P–Q), and include derived human-biased response patterns across selected genes (R). **(S–U)** At 54 hrs, divergent BMP7 responses are enriched for morphogenetic and cytoskeletal processes. Divergent rGenes are associated with forebrain development, epithelial-to-mesenchymal transition (S–T) and actin cytoskeleton and morphogenesis-related pathways (U).

## SUPPLEMENTAL INFORMATION

**Table S1: PSC line and directed differentiation reagent metadata.**

**Table S2: Morphogen screen chemical metadata and viability screen results.**

**Table S3: scRNA-seq cell class, cell type, and cell state markers for dataset subsets**

**Table S4: Dreamlet stimulus response differential gene expression results and species comparison**

**Table S5: Energy distance results Table S6: rGene correlation results**

**Table S7: Milo differential cellular abundance results Table S8: rGene and conserved rGene enrichment results**

**Table S9: Dreamlet baseline and response context species differential gene expression and context comparison**

**Table S10: Divergent baseline gene and divergent rGene enrichment results Table S11: Description of datasets for genomic enrichment analyses.**

**Table S12: Human-specific genomic features for genomic enrichment analyses.**

## METHODS

### Maintenance of great ape PSC lines

Hominid PSCs were cultured in PSC Media + IWR1 (2 µM) on Growth Factor-Reduced Matrigel (Corning) (Table S1) as in Schaefer et. al.^77^ When cultures approached or reached confluence, cells were cluster passaged with Cell Release solution (30 mM NaCl and 0.5 mM EDTA in PBS) and plated in PSC Media + IWR1 (2 µM) + CEPT^121^ for a minimum of one hour before removing CEPT. PSC lines were passaged 3-6 times prior to directed differentiation to synchronize their growth. When thawing cryopreserved cultures, CEPT was added overnight to improve culture survival. PSC cultures were cryopreserved in Bambanker (Bulldog Bio).

### Directed differentiation of telNECs from great ape PSC lines

On Day -1, adherent PSC cultures were washed with Cell Release and then dissociated with Cell Release, testing for dissociation in 5 min increments and gently triturating the cells until small clusters of several to dozens of cells were obtained using wide orifice pipette or serological pipette tips. Cell suspensions were diluted with two volumes of PSC media + CEPT (without IWR1), counted by packed cell volume (PCV) in duplicate, and seeded at 50,000 or 100,000 cells/cm^2^ onto growth factor-reduced matrigel. Media changes during Day 0-7 were performed according to Table S1, with media changes performed each day from Day 4-7 if media acidification was observed. At Day 7, cultures were washed with Cell Release and dissociated with room temperature Accutase, typically for 10 mins. After removal of Accutase, cultures were firmly tapped to dislodge adherent cells before addition of Cell Release for gentle trituration to a single cell suspension. Two to three volumes of Day7+ Media + CEPT were added to the cell suspension before counting with a Cellaca PLX using a 1:1 dilution of cell suspension to Viastain AOPI Staining Solution and optimized custom imaging parameters. Cultures were seeded at 650,000 cells/cm^2^ on Polyethylenimine (PEI)-treated and growth factor-reduced matrigel-coated plates. At Day 14, cultures were washed with Cell Release and dissociated with Cell Release into clusters of several to dozens of cells, typically for 15 mins. Two to three volumes of Day7+ Media + CEPT were added to the cell suspension before counting as in the Day 7 passaging. Cultures were seeded in Day7+ Media + CEPT at a 1:3-1:6 passaging ratio to normalize growth between individual cultures before removal of CEPT after one hour. At each passage, cultures were seeded in parallel for quality control ICC on CellVis glass bottom 96-well plates. During culture passaging, an equal number of human and non-human great ape individual cultures were divided amongst technicians RCM and BJP with individual identities semi-randomly assigned to each technician to control for technician handling differences. Prior to cryopreservation, differentiated cultures were treated with Day7+ Media + CEPT for at least one hour. Cultures were passaged and processed as at Day 14, resuspending in Bambanker and cooling to -80°C in a controlled-rate cooler before transferring to liquid nitrogen. For additional details regarding the differentiation protocol, please refer to Fig. S1 and Table S1.

### TelNEC expansion, intraspecies pooling, and interspecies pooling

Cryopreserved stocks of PSC-derived telencephalic NEC cultures from 4 different differentiation batches banked on Day 13, 14, 18, or 21 were thawed in stages into Day 7+ Media + CEPT so as to synchronize their chronological age. All cultures from the same differentiation/cryopreservation batch were treated identically leading up to passage synchronization to control for batch effects. At thawing and at each passage, cultures were seeded at two to four densities using a twofold or threefold dilution series to ensure culture survival and growth. When cultures approached or reached confluence, cells were passaged with Cell Release solution and plated in Day 7+ Media + CEPT. At Day 22, 20 ng/mL of FGF2-G3 was added to Day 7+ Media. After Day 25, CEPT was added to media only overnight following a passage. At Day 33, all cultures from all differentiation batches were synchronously passaged and treated identically thereafter. At Day 35, human and chimpanzee cultures from six individuals per species and one orangutan culture were selected based on quality control and batch and sex matching within differentiation batches. Cultures were passaged with Cell Release, counted with a Cellaca PLX using a 1:1 dilution of cell suspension to Viastain AOPI Staining Solution and custom imaging parameters, and pooled in equal numbers of live cells into intraspecies pools. At Day 39 and 43, intraspecies human or chimpanzee cultures and the orangutan culture were passaged. At Day 46, FGF2-G3 was replaced with FGF2 at 20 ng/mL. At Day 48, intraspecies pools were passaged with Accutase and pooled into interspecies pools, targeting equal numbers of FOXG1+/NKX2-1+ cells per species, with orangutan cells added with a 25% increase relative to the expected single individual representation. Pools were plated at 25,000 cells per cm^2^ in Differentiation Media + CEPT in 24- or 96-well glass-bottom plates on Growth Factor-Reduced Matrigel. At Day 49, media was changed to Differentiation Media, and cultures were fed on Day 50 and 53, as well. Starting at Day 54, interspecies pools were treated with different morphogen perturbations or controls every day for 7 days.

### Immunocytochemistry, imaging, and image analysis

Samples of adherent cell cultures in 96-well glass-bottom plates were washed once with 100 µL of DMEM/F12 1:1 and then fixed with 2% paraformaldehyde (PFA) in DMEM/F12 1:1 (i.e. 100 µL of 4% PFA + 100 µL DMEM/F12 1:1) for 15 minutes at room temperature. Then, samples were washed with 100 µL of phosphate-buffered saline (PBS) twice followed by fixation/permeabilization with ice-cold 90% methanol for 10 minutes at room temperature. Again, samples were washed twice with 100 µL PBS before storing in PBS with 0.1% sodium azide. For primary antibody staining, antibodies were diluted to working concentration in 100 µL of Blocking Buffer, composed of 0.3% Triton X-100 and 5% bovine serum albumin (BSA) in PBS, and incubated for 1 hour at room temperature. Samples were washed twice with Wash Buffer, composed of 0.1% Tween-20 in PBS. For secondary antibody staining, antibodies were diluted to working concentration in 100 µL of Blocking Buffer along with 1 µg/mL of DAPI and incubated for 1 hour at room temperature while protected from light. Finally, samples were washed twice with 100 µL of Wash Buffer and stored in PBS with 0.1% sodium azide.

For each sample, 3-4 images were acquired with a EVOS M7000 Imaging System microscope using a 10X objective. Segmentation of cells was performed with either CellProfiler version 4.2.5 or cellpose version 3.0.10.

In the CellProfiler workflow, objects for each channel were segmented using IdentifyPrimaryObjects with the following parameters: threshold strategy = global, thresholding method = minimum Cross-Entropy, and typical pixel diameter of objects = 9-20. For each marker and channel, threshold correction factor as well as lower and upper bounds on threshold parameters were determined heuristically for defining positive and negative objects. Counts of single-, double-, and triple-positive objects as well as total objects were then computed per sample using RelateObjects and FilterObjects, and percentages of different cell populations were quantified using CalculateMath.

In the cellpose workflow, DAPI+ objects were segmented using the ‘cyto3’ model with or without ‘deblur’, with ‘flow threshold’ and ‘cellprob threshold’ parameters determined heuristically. Per-object mean fluorescence intensity was computed using skimage.measure function regionprops_table, and intensity thresholds defining positive and negative objects were determined heuristically. Counts of single-, double-, and triple-positive objects as well as total objects were then computed. For the stimulus response analysis, a binomial generalized linear mixed-effects model was fit with the glmer() function using a logit link to model the proportion of positive cells and using the ‘bobyqa’ optimizer. Fixed effects included species, condition, and their interaction, while biological replicate was included as a random intercept to account for repeated measures within biological replicates and inter-replicate variability. Estimated marginal means were computed for stimulus treatment within each species. Stimulus effects were quantified by contrasts comparing each condition to the vehicle control within species. Contrast estimates were reported on the log-odds scale and converted to odds ratios with corresponding 95% confidence intervals by exponentiation.

To compare changes in culture density, the number of DAPI-positive objects (n₀) from cellpose segmentation was calculated per image from three biological replicates (i.e. wells) and three to four technical replicate images (i.e. fields of view). To test for effects of species and experimental condition while accounting for biological replication, n₀ was modeled with a generalized linear mixed-effects model with a negative binomial distribution using the glmmTMB package function glmmTMB with family = nbinom2(). Fixed effects included species, condition, and their interaction, with a random intercept used for biological replicate, according to the following model form:

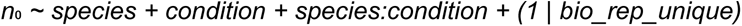

Estimated marginal means were computed for each condition separately in each species on the log scale. To compare experimental conditions to vehicle within each species, we used emmeans function contrast() with method = “trt.vs.ctrl”, ref = “Vechicle”, and by = “species”. Contrasts on the log scale were transformed to obtain fold-changes in n₀ relative to vehicle with 95% confidence intervals and associated p-values using emmeans package function summary() with type = “response” and infer = c(TRUE, TRUE). Statistical significance was assessed using Wald tests. Obtained p-values were adjusted for multiple testing using the Benjamini-Hochberg FDR procedure with p.adjust() function with method = “fdr”.

### Chemical viability screening

To assess culture viability dose responses to chemicals over time, multi-individual intraspecies cultures of chimpanzee telNECs were cultured on 96-well plates and treated with various chemical conditions in triplicate or quadruplicate. After 2 days and 9 days, media was removed from cultures and replaced with 90 µL DMEM/F12 1:1 and 10 µL of 10X PrestoBlue HS Cell Viability Reagent (ThermoFisher Scientific) and calculated flux was quantified every 30 mins until the fluorescence signal began to saturate. Calculated flux values correspond to the slope of the fluorescent signal over time as computed by the Promega plate reader software. Average calculated flux was measured across DMSO vehicle wells for each 96-well plate. Calculated flux for all other conditions were normalized to the same-plate vehicle average. One-sample, two-sided t-tests were then performed for each condition to assess whether the distribution of normalized flux values differed from the vehicle mean. To account for multiple hypothesis testing across conditions, p-values were adjusted using the Benjamini-Hochberg FDR procedure with a significance threshold of FDR < 0.10.

### Sample processing, MULTI-seq barcoding, and scRNA-seq library preparation

To prepare adherent cultures for MULTI-seq barcoding, cultures were dissociated in Accutase for 5 mins, diluted in three volumes of cold Cell Release, and centrifuged at 500 xg at 4°C for 5 mins. All following centrifugation steps were performed identically. After removal of the supernatant, cells were resuspended in a normalized volume and counted using the Cellaca PLX (Revvity) after a 1:1 dilution in Viastain solution (Revvity). Cell suspension volumes for 1.5e6 cells (i.e. three 0.5e6 MULTI-seq reactions) were transferred to a 96-well round bottom plate. Cell pellets were centrifuged and washed three times in 200 µL of cold PBS-EDTA before resuspending in 300 uL of PBS (i.e. 100 µL per MULTI-seq reaction). Volume of cell suspension for one MULTI-seq reaction (i.e. 100 µL) was added to 22 µL of 10 uM MULTI-seq Anchor + 10 uM MULTI-seq Barcode in PBS, incubated on ice for 5 mins, transferred in full to 22 uL of 10 uM MULTI-seq Co-Anchor in PBS, incubated on ice for 5 mins, transferred in full to a 96-well round-bottom plate, and washed with an equal volume (i.e. 144 µL) of cold 5% BSA in PBS. Cell pellets were centrifuged and washed three times with 200 µL of cold 2% BSA.

Samples were divided into 4 groups, and equal volumes of each sample were pooled in 4 different pools. Excess cells from media and vehicle controls were spiked into these pools to increase their representation twofold. The same was done for the LY411575 condition only for the 54 hr and 7 day time points. Sample pools were filtered through a 75 µm filter (Flowmi). Cell counts from the sample pools were used to pool equal numbers of cells from each pool in two iterative pooling steps. Cell counts from the final pool were used to resuspend the final pool to a target concentration of 6e6 cell/mL in cold 1% BSA in PBS after centrifugation and aspiration of the supernatant. The final pool was filtered through a 30 µm filter (Flowmi), counted, and resuspended in cold 1% BSA in PBS for a target final concentration of 2e6 cells/mL. GEM generation and barcoding were performed according to 10X Genomics User Guide CG000416 | Rev D, with a targeted capture of 115,400 cells.

### Library preparation and sequencing

To prepare gene expression libraries, post GEM-RT cleanup, cDNA amplification, and 3’ gene expression library construction were performed according to 10X Genomics User Guide CG000416 | Rev D, with 11 and 12 PCR cycles used for the cDNA Amplification and Sample Index PCR steps, respectively. The MULTI-seq additive primer was added during cDNA amplification at Step 2.2a. The quality of the resulting libraries was assessed using the 4200 Tapestation using High Sensitivity D1000 ScreenTape/Reagents (Agilent) before they were sequenced on the Illumina NovaSeqX 25B platform with the recommended cycle parameters targeting 25-30k mean reads per cell.

MULTI-seq barcode libraries were prepared according to McGinnis et. al.^122^ The MULTI-seq barcode fraction was saved during cDNA cleanup Step 2.3d. The library preparation PCR was performed with 10 cycles. MULTI-seq barcode libraries were sequenced along with the gene expression libraries, targeting 1k mean MULTI-seq barcode read counts per cell.

### Single-cell RNA-seq data processing

We aligned single-cell RNA-seq data using CellRanger version 6.1.2, with the option –create-bam set to true. The reference genome was the imputed common ancestor of humans, chimpanzees, and bonobos, produced along with a homology-based annotation as part of a prior study,^123^ as described previously.^77^ Sequencing and alignment yielded per cell measures of 27,300 mean reads, 7,010 median counts, 3,128 median genes, and 944 median MULTI-seq counts. Datasets were concatenated with anndata package function concat using an outer join.

### Demultiplexing stimulus, species, and individual from sequencing reads

We demultiplexed stimulus, species, and individual identities from the sequencing reads using version 0.1.0 of the CellBouncer suite of tools^77^. To assign stimulus identity to cells based on MULTI-seq barcode counts, we used the demux_tags program with the filtered list of cell barcodes (i.e. following preprocessing and quality control) as inputs with default parameters.

To assign individual and species identity, first produced a set of variants using whole-genome sequencing data from our cell lines that were collected as part of a prior study^77^. Whole-genome sequencing reads were aligned to the human/chimpanzee/bonobo ancestor genome using minimap2^124^, name-sorted with samtools^125^ sort, modified to add mate pair information with samtools fixmate, position-sorted with samtools sort, and duplicate-marked using samtools markdup. Variants were then called using FreeBayes^126^ with default parameters and filtered to high-quality biallelic sites where no more than 50% of individuals are missing a genotype, via bcftools^125^ -m 2 -M 2 -q 100 -i “F_MISSING < 0.5”. To increase computational efficiency, we then downsampled variant sites so that no more than 10 million variants representing fixed differences on a given phylogenetic branch were kept. After inferring each cell’s individual of origin, we inferred species identity based on the individual identity.

As a second measure of species identity, we used CellBouncer’s demux_species program. When building reference k-mer lists, we used k = 32, sampled all available k-mers, and filtered for orthologous transcripts (options -k 32 -N -1 -O to the program demux_species_ref.py). We extracted transcript sequences using gffread^127^ version 0.12.6 with the command gffread -F -w [output_fasta] -g [input_fasta] [input_annotation], using the human genome build hg38 and the GENCODE v36 [PMID 39565199] GTF-format annotation, and with chimpanzee and orangutan genome builds and annotations taken from a hierarchical alignment (HAL) of current best assemblies produced and annotated as part of two recent studies [PMID 33953399, PMID 29880660]. As an extra measure of quality control, we excluded cells for which the species identity inferred this way disagreed with the species identity inferred from individual of origin.

### Ambient RNA quantification and removal

We quantified ambient RNA using CellBouncer’s quant_contam program and the cell-to-individual assignments from demux_vcf (see Demultiplexing stimulus, species, and individual from sequencing reads). This program infers ambient RNA using the rates at which cells mismatch their expected (true) genotypes, once their individuals of origin are identified; ambient RNA is modeled as a weighted mixture of all genotypes present in the pool. Since we aligned all species in our data set to a common reference genome, cross-species ambient RNA contamination was handled appropriately. After inferring the ambient RNA profile and per-cell contamination rates, we also used quant_contam to produce decontaminated gene expression matrices, using Leiden clusters inferred from the initial gene expression matrices (see scRNA-seq data preprocessing, clustering, integration, and annotation) to model endogenous gene expression profiles. These decontaminated expression matrices were used in all downstream analyses.

### scRNA-seq data preprocessing, clustering, integration, and annotation

Analysis of scRNA-seq data was performed using scanpy^78^ with AnnData structures.^128^ All datasets were concatenated and filtered to genes with more than 10 cells and to cells with more than 100 genes, fewer than 20% mitochondrial reads, and fewer than 10% ribosomal reads. The verteporfin perturbation was removed on the basis of higher mitochondrial and ribosomal read percentages. Time point, perturbation, species, and individual were added to the AnnData object and doublets for the latter three variables were removed. Following standard normalizing and logarithmizing of counts as well as dimensional reduction and leiden clustering, we annotated transcriptomic clusters as mesenchyme, midbrain and hindbrain, or telencephalon regions. We subset the dataset to the telencephalic cells and re-performed principal component analysis (PCA). As species formed distinct clusters unrelated to regional or cell class identity, we performed dataset integration by correcting the PCA dimensions using the scanpy external package function harmony_integrate across species, time point, and individual with the following parameters: theta = [2,2,2], lamb = [1,1,1], and sigma = 0.1.

Following clustering, clusters were annotated according to their cell class based on visualization of canonical cell class markers as well as their identification within top marker gene rankings from scanpy function rank_gene_groups() using the wilcoxon method. We annotated *SOX2*+/*HES1*+/*NES*+ clusters as NECs, *GADD45G*+/*HES6*+/*DLL3*+ clusters as IPCs, and *GAP43*+/*DCX*+/*ELAVL3*+ cells as immNs. We distinguished excitatory or inhibitory IPCs based on expression of *NEUROG1, NEUROD4*, and *NHLH1* or *ASCL1*, *DLX2*, and *DLX5*, respectively. We distinguished excitatory or inhibitory neurons based on expression of *SLC17A6* and *SLC17A7* or *GAD1*/2 and *SLC32A1*, respectively. Cell-cycle phase was assigned using scanpy function score_genes_cell_cycle() with curated S-phase genes (*MCM5, PCNA, TYMS, FEN1, MCM2, MCM4, RRM1, UNG, GINS2, MCM6, CDCA7, DTL, PRIM1, UHRF1, HELLS, RFC2, RPA2, NASP, RAD51AP1, GMNN, WDR76, SLBP, CCNE2, UBR7, POLD3, MSH2, ATAD2, RAD51, RRM2, CDC45, CDC6, EXO1, TIPIN, DSCC1, BLM, CASP8AP2, USP1, CLSPN, POLA1, CHAF1B, BRIP1, E2F8*) and G2/M-phase genes (*HMGB2, CDK1, NUSAP1, UBE2C, BIRC5, TPX2, TOP2A, NDC80, CKS2, NUF2, CKS1B, MKI67, TMPO, CENPF, TACC3, SMC4, CCNB2, CKAP2L, CKAP2, AURKB, BUB1, KIF11, ANP32E, TUBB4B, GTSE1, KIF20B, HJURP, CDCA3, CDC20, TTK, CDC25C, KIF2C, RANGAP, NCAPD2, DLGAP5, CDCA2, CDCA8, ECT2, KIF23, HMMR, AURKA, PSRC1, ANLN, LBR, CKAP5, CENPE, CTCF, NEK2, G2E3, GAS2L3, CBX5, CENPA*).^129^ Lastly, one cluster was composed of lower quality cells escaping the initial thresholding and was therefore removed from the dataset.

To examine cell type heterogeneity within the progenitor cell classes not relating to other biological or technical variables, we subset the dataset to the NEC and RG classes and performed dataset integration with scVI.^80^ Specifically, we set up the scVI model to integrate species while considering perturbation, time point, individual, sex, sequencing batch, differentiation batch, and cell-cycle phase as categorical covariates using the following parameters: use_layer_norm = “both”, use_batch_norm = “none”, encode_covariates = True, dropout_rate = 0.2, and n_layers = 2. We trained the scVI model using the following parameters: early_stopping = True, max_epochs = 100, train_size = 0.75, check_val_every_n_epoch = 5. After neighbor finding, dimensional reduction, and leiden clustering, we identified two low quality clusters that were removed from the dataset. We identified a gradient of gene expression changes across NECs which segregated into 4 broad clusters, which we labeled as dorsal NEC, dorsal/ventral NEC, ventral NEC and vRG. Dorsal NECs were initially annotated on the basis of *PAX6* and *GLI2* expression, principally, while ventral NECs were initiallyannotated on the basis of *NKX2-1, PROX1*, *LMO1*, and *FRZB* expression, with both annotations supported by mappings to a macaquer reference *in vivo dataset*.^84^ We note that many dorsal NECs displayed increased expression of early NEC genes *SOX1* and *LRRN1* while most ventral NECs displayed increased expression of *EPHA5*, *PTN*, and *NFIX*, supporting an alternative annotation of early versus late NEC fate and that these interpretations are not mutually exclusive. Lastly, we note that the marker gene expression of ventral NECs is consistent with a developmental stage prior to specification of the distinct ganglionic eminences.

To examine transcriptional heterogeneity within NECs relating to perturbations but not other biological or technical variables, we subset the dataset to NECs and performed dataset integration with scVI. Again, we set up the model to integrate species while considering the aforementioned study variables excluding perturbation and using the same scVI model setup and training parameters. After neighbor finding, dimensional reduction, and leiden clustering, we identified one cluster whose top ranked marker genes included several canonical BMP response genes, which we annotated as NEC_BMP_response. Similarly, we identified a second cluster whose top ranked marker genes included several canonical FGF and ERBB response genes, which we annotated as NEC_FGF_ERBB_response.

To explore cell type heterogeneity within differentiated cell classes not relating to other biological or technical variables, we subset the dataset to the IPC and immN classes. We performed dataset integration with scVI in the same manner as the progenitor cell classes, adding percent counts mitochondrial reads, percent counts ribosomal reads, number of genes by counts, and total counts as continuous covariates to the model. After neighbor finding, dimensional reduction, and leiden clustering, we annotated clusters based on marker genes from the literature, cluster markers from a macaque reference *in vivo* dataset,^84^ and mappings to this reference. Within inNs, we identified transcriptomic clusters of ventral (*NKX2-1*, *LHX6*, *LHX8*, *SHH*, *GBX1*), ventromedial (*NKX2-1*, *HMX3*, *POU6F2*), and GnRH neuronal (*GNRH1*, *SP8*, *MEIS2*) identities (Fig. 1, Fig. S4H-J, Table S3). Among exNs, we identified clusters expressing markers of preplate neurons (*TMEM163*+/*EBF2*+), including immature subplate neurons and Cajal-Retzius neurons (*LHX5*+/*TLE4*+). ^81–83,130^ We annotated these population as of presumptive pallial or subpallial/pallial-subpallial boundary origin, distinguished by co-expression of *EOMES*, *TBR1*, and *RSPO3* or *NKX2-1*, *SIM1*, and *OLIG1/2* or along their respective developmental trajectories (Fig. 1N, Fig. S4H-J). However, without markers of mature neurons, we were unable to resolve subplate versus Cajal Retzius neuron identity.

### Semi-supervised probabilistic annotation based on macaque *in vivo* reference atlas

Given that we observed a rare vRG-like cell type, no cluster displaying a clear outer RG signature, neurogenesis of preplate but not upper layer exNs, and no astrogenesis, we reasoned that the cultures likely represented a first trimester developmental stage. Therefore, we subset the macaque reference dataset to macaque ages corresponding to the first trimester of human gestation.^131^ We rebalanced the reference dataset across developmental stages and cell subtypes by subsetting to a maximum of 1,000 cells for each combination of stage and cell subtype. We annotated cell types based on groups of cell subtypes. The current study query dataset was rebalanced across species and cell type/state and subset to 1,000 NECs or 500 cells for other cell types per combination of species and cell type/state. We preprocessed the query and reference dataset identically, dropping cells with fewer than 100 genes, dropping genes with fewer than 10 cells, and normalizing and logarithmizing counts. Both datasets were concatenated and subset to shared genes before again normalizing and logarithmizing counts. The top 2,000 highly variable genes were determined from the reference dataset subset, and the concatenated dataset was subset to these features. The dataset was then integrated with scVI using a batch key representing different sample groups in each dataset and a categorical covariate key representing the different individuals in each dataset. The scVI model was set up with the following parameters: use_layer_norm = “both”, use_batch_norm = “none”, encode_covariates = True, dropout_rate = 0.2, n_layers = 2. The scVI model was trained with the following parameters: early_stopping = True, max_epochs = 300, train_size = 0.75, check_val_every_n_epoch = 10. To perform probabilistic annotation, we set up a scANVI model with the trained scVI model and cell type annotations as inputs. We fit the scANVI model with the following parameters: max_epochs = 300 and n_samples_per_label = 100. Cell type label probabilities were predicted using the predict function, with 96.14% accuracy for the predicted reference labels. Query cell types/states were annotated based on majority vote of cells within that cell type/state. The latent representation from the scANVI model was acquired with the get_latent_representation function, and a k-nearest-neighbor graph (k = 100) was reconstructed using Euclidean distances in the scANVI latent embedding. The resulting directed connectivity matrix was restricted to edges pointing from all cells to reference cells with their associated ages. For each cell, we then calculated the weighted mean age of its reference neighbors by multiplying connectivity weights by reference ages and normalizing by the total connectivity weight.

### Pseudotime analysis

To order telencephalic cells from each species along a pseudotime, we first converted our dataset to a monocle object using monocle package function new_cell_dataset with cell metadata, gene metadata, and the cell by gene expression matrix as inputs. We clustered cells with monocle function cluster cells with the UMAP reduction method and randomly subset the dataset to up to 10,000 cells per species. We defined ‘root’ cells as dorsal NECs that were untreated or treated with vehicle treatments at the 6 hr time point and ordered cells along pseudotime using the monocle function order_cells().

### Differential gene expression analysis

We performed differential gene expression analysis using the dreamlet package,^87^ which uses a pseudobulk approach with a precision-weighted linear mixed effects model. First, we defined sample group as the combination of species, time point, and perturbation and sample as the combination of sample group and individual. Second, we pseudobulked the scRNA-seq count data by cell class. Then, we ran voom-style normalization for pseudobulk counts within each cell class using processAssays with the model *∼ (1|species) + (1|perturbation) + (1|time_point)* and the following parameters: min.count = 5, min.cells = 40, min.samples = 4, and min.prop = 0.0178. To determine which variables to include in the differential gene expression model, we first performed a variancePartition analysis to assess the fraction of expression variation explained by each variable. We used the model *∼ (1|species) + (1|perturbation) + (1|time_point) + (1|individual) + (1|seq_batch) + (1|sex*) for the fitVarPart function, electing to drop seq_batch and sex from further analysis as they explained a lesser degree of expression variation. Finally, we modeled differential gene expression using dreamlet with a means model for factors of the form ∼ 0 + sample_group + (1|individual), specifying individual as a random effect and using the result of processAssays and default parameters as inputs. We defined contrasts for testing stimulus response with the form:

*sample_groupSpecies_Perturbation_TimePoint - sample_groupSpecies_Vehicle_TimePoint*

We defined contrasts for testing species divergence in stimulus response with the form:

*(sample_groupHuman_Perturbation_TimePoint - sample_groupHuman_Vehicle_TimePoint) - (sample_groupChimp_Perturbation_TimePoint - sample_groupChimp_Vehicle_TimePoint)*

We defined contrasts for testing species divergence in the baseline context with the form:

*(sample_groupHuman_Untreated_TimePoint + sample_groupHuman_Vehicle1_TimePoint + sample_groupHuman_Vehicle3_TimePoint) / 3 - (sample_groupChimp_Untreated_TimePoint + sample_groupChimp_Vehicle1_TimePoint + sample_groupChimp_Vehicle3_TimePoint) / 3*

We defined contrasts for testing species divergence in the response context with the form:

*sample_groupHuman_Perturbation_TimePoint - sample_groupChimp_Perturbation_TimePoint*

We defined divergent rGene categories with the following algorithm. Baseline only genes were significant for species differences at baseline but not in response. Response only genes were significant for species differences in response but not at baseline. Enhancing or reversing genes were significant for species difference in response and baseline, with greater or lesser absolute value of divergence in response, respectively. Reversing genes were significant in baseline and in response with opposing signs of divergence direction.

We defined conserved rGenes as rGenes with showed significant gene expression responses and conserved with the same direction of effect for a stimulus in both human and chimpanzee (FDR < 0.10), and divergent rGenes as rGenes with significant divergence in their response to stimuli between these species (FDR < 0.10, Methods). Divergent rGenes showed significant species differences in response. by stimulus combinations modeled that could not be categorized as conserved or divergent but responded significantly in at least one species were annotated as ‘other rGenes’. Finally, genes that did not reach significance in either species were annotated as ‘non-rGenes’.

### Energy distance analysis

To quantify stimulus responses across cell types in a single metric, we calculated the energy distance (E-distance) between stimulus groups (i.e. cells belonging to each stimulus), sample metagroups (i.e. cells belonging to each combination of stimulus and species), and sample groups (i.e. cells belonging to each combination of stimulus, species, and time point) and their corresponding vehicle controls using the pertpy package function tools.DistanceTest() with metric = “edistance”, n_perms = 20,000, and obsm_key = ‘X_PCA_harmony’. Pairwise E-distances were similarly calculated with the pertpy package function tools.Distance() with pt.tl.Distance with metric = “edistance” and obsm_key = “X_pca_harmony”.

### Correlation of rGene changes across sample groups

To quantify correlation in response gene changes across sample groups, we first subset the dreamlet differential gene expression results to only rGenes. Then, we calculated the pairwise Pearson’s correlation coefficients between sample groups using the base R stats package function cor() with use = “pairwise.complete.obs” and method = “pearson”.

### Differential cellular abundance analysis

To test for differential cellular abundance in response to stimuli, we used the miloR^89^. First, we constructed a KNN graph with the buildGraph() function using parameters k = 500 and d = 20, with dimensions from harmony-corrected PCA. We then built cellular neighborhoods using the makeNhoods() functions with parameters prop = 0.005, k = 500, and d = 20 following a parameter sweep for optimization of prop and k such that multiple neighborhoods populated each cell type and were distributed densely across the neurogenic trajectory. Third, we counted cells of each sample group in each neighborhood using the countCells() function. Then we calculated the distance between neighborhoods for Spatial FDR correction using the calcNhoodDistance() function. Finally, we tested differential abundance using the testNhoods() function, defining contrasts for stimulus response and species divergence in stimulus response as in the differential gene expression analysis. For visualization, neighborhoods were assigned to a cell type by majority vote of resident cells.

### Gene set over-representation and enrichment analysis

For the enrichment tests performed on the rGene categories as a whole, we tested whether genes (which passed the model criteria for dreamlet) from one category were enriched in that gene set relative to genes from a second category for each gene set using a 2×2 contingency table of in-gene set versus not-in-gene set membership. Enrichment was assessed with a two-sided Fisher’s exact test, and odds ratios and 95% confidence intervals were reported. Obtained nominal p-values were corrected for multiple hypothesis testing using the Benjamini-Hochberg FDR procedure across gene sets, and gene sets with fewer than 100 genes were excluded from the final results. The set of eGenes was obtained from GTEx eQTL Analysis v10 (https://www.gtexportal.org/home/downloads/adult-gtex/qtl), filtering genes with a q-value < 0.05. The set of GWAS genes and LoF-intolerant were obtained from the MacArthur Lab (https://github.com/macarthur-lab/gene_lists), with the LoF-intolerant gene set as the combination of the essential genes in mice, shRNA perturbation, CRISPR perturbation, or autosomal recessive genes from Online Mendelian Inheritance in Man.

For the over-representation analysis for GO:BP terms within convergent and divergent rGenes for each perturbation, we used the clusterProfiler package^132^ function enricher() with the following parameters: pAdjustMethod = “fdr”, pvalueCutoff = 0.05, qvalueCutoff = 0.10, minGSSize = 5 or 10, and maxGSSize = 500. For the over-representation analysis performed on convergent and divergent rGenes for each perturbation, we used clusterProfiler package function gseGO() with the following parameters: ont = “BP”, minGSSize = 5, eps = 0, and by = “fgsea”. The results were filtered for adjusted p-values less than 0.10.

For the over-representation analysis for disease-associated genes within convergent and divergent rGenes for each perturbation, we using the disgenet2r package^133^ function disease_enrichment() with the following parameters: common_entities = 3, max_pvalue = 0.10, vocabulary = ’HGNC’, and database = ’CURATED’. The results were filtered for adjusted p-values less than 0.10.

### Bulk RT-qPCR and stimulus response validation and analysis

Multi-individual, intraspecies telencephalic cultures of human and chimpanzee were cultured in 24-well plates and treated with vehicle, BMP7 (10 ng/mL), BMP7 (100 ng/mL), or LDN193189 (0.5 µM) for 54 hrs. Samples were washed once with 350 µL of PBS and lysed with 350 µL of Qiagen RLT RNeasy Lysis Buffer. RNA was extracted using the Qiagen RNeasy Mini Kit. cDNA was synthesized using the Maxima First Strand cDNA Synthesis kit from 1 µg of RNA with the following thermocycler parameters: 25°C for 10 min, 50°C for 15 min, 85°C for 5 min. RT-qPCR reactions were set up using 384-well plates with 10 µL reaction volumes containing 1X PowerUp SYBR Green Master Mix, 10 ng of cDNA, and 0.25 µM of forward and reverse primers. No template controls (NTCs) were amplified using each unique primer pair and no reverse transcriptase controls (NRTs) were amplified with the endogenous housekeeping control GAPDH primer pair. RT-qPCR was performed on a QuantStudio 5 Real-Time PCR System (Applied Biosystems) using the Relative Quantification module in Fast mode with a heated cover at 105°C. Cycling included a hold stage (50°C for 2 min, 95°C for 2 min), followed by 40 cycles (95°C for 15 s, 55°C for 20 s, 60°C for 30 s) with fluorescence collected at each step. Melt curve analysis (95°C for 15 s, 55°C for 20 s, 95°C for 15 s) was performed with continuous acquisition. All reactions were quality controlled based on NTCs, NRTs, and single expected product melting temperatures. RQ values were obtained relative to the GAPDH housekeeping gene using the ThermoFisher Cloud software RQ tool with vehicle-treated human cultures set as the reference level.

For each gene, RQ values were log2 transformed and fit to a linear model with species, treatment, and their interaction using the lm() function with the following model form:

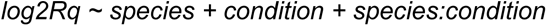

Type III sums-of-squares ANOVA was used to test these main effects and their interaction using the car package Anova() function with type = 3. Estimated marginal means were computed for all species by treatment combinations with emmeans package function emmeans. Within each species, responses were quantified as contrasts of each condition versus the vehicle control using the emmeans package function contrast() with method=”trt.vs.ctrl”, by = “species”, and ref = “vehicle”. Statistics were performed with Type III Wald F-tests and p-values and 95% confidence intervals were reported using emmeans function summary() with infer = TRUE. To assess divergent responses between species, the within-species response contrasts were then compared across species using the emmeans package function contrast() with method=”pariwise” and by=”contrast”. Statistics were performed with Wald t-tests on linear contrasts of model coefficients and p-values and 95% confidence intervals were reported using emmeans function summary() with infer = TRUE. Nominal p-values were considered as significant if less than 0.10.

### Human-specific genomic feature class enrichments

We assigned human-specific genomic features to overlapping or proximal genes for downstream enrichment analysis. First, we lifted *de novo* gene features from hg19 to hg38 and hDel features from panTro6 to hg38 (after adding 500 bp 3’ and 5’ of hDel features to facilitate liftover of elements absent from the human genome by definition), retaining the largest mapped segment in the case of multimapping. Human gene, exon, and intron annotations were obtained from Ensembl (EnsDb.Hsapiens.v86) and standardized to the hg38 reference with UCSC-style chromosome naming, retaining only canonical chromosomes. Each lifted genomic feature set was intersected with these annotations to classify intervals as exonic, intronic, or intergenic using a hierarchical priority scheme (exonic > intronic > intergenic). Exonic and intronic features were assigned overlapping gene symbols based on direct interval overlap, while intergenic features were annotated with the nearest upstream and downstream genes and their genomic distances. Rare non-exonic hSegDups or *de novo* genes were dropped. Overall, for intergenic features, 75% fell within 125.0 kb of a gene, with a median genomic distance of 48.2 kb. After filtering intergenic features for those within 500 kb of a gene, we found that we retained more than 98.6% of features across all classes. Further, 85.4% of hIns, 85.5% of hDels, 84.2% of hCONDELs, 88.1% of HAQERs, 89.8% of VNTRs, and 78.2% of zooHARs fell within 100 kb of a gene.

## ACKNOWLEDGEMENTS

We thank Danny Conrad (Gartner Lab, UCSF) for providing MULTI-seq reagents and protocols; Jenelle Wallace for scripts, guidance on differential gene expression analysis, and feedback on the manuscript; Jingwen Ding for scripts and input on differential cellular abundance analysis; Gaurav Rathore for scripts and input on dataset integration, Enakshi Sinniah for scripts and guidance on gene set enrichment analysis, and members of the Pollen lab for helpful discussions and feedback throughout the project. Sequencing was performed at the UCSF Center for Advanced Technology (CAT), supported by UCSF PBBR, RRP IMIA, and NIH grant 1S10OD028511-01.This work was supported by a Weill Neurohub Fellowship (N.K.S.); National Institutes of Health grants R01AG087959, R01MH134981, and DP2MH122400 (A.A.P.); NSF grant #2134955 (A.A.P.); the Pershing Square Foundation; Schmidt Futures; the Shurl and Kay Curci Foundation; and an Innovative Genomics Institute Award (A.A.P.). A.A.P. is a New York Stem Cell Foundation–Robertson Investigator. This project was also supported in part by the Emory National Primate Research Center (ORIP/OD P51OD011132).

## AUTHOR CONTRIBUTIONS

- RCM, BJP, and AAP conceived of the project and designed the experiments.
- RCM and AAP supervised the experiments and analysis.
- RCM, BJP, and DMS performed stem cell culture and directed differentiations.
- RCM and BJP performed the morphogen perturbation scRNA-seq experiment.
- RCM performed gene expression and MULTI-seq barcode library preparation.
- RCM performed scRNA-seq data preprocessing, clustering, annotation, and reference mapping.
- RCM performed differential gene expression analysis
- RCM and DSA performed differential cellular abundance analysis
- RCM and NKS performed demultiplexing of perturbation, species, and individual
- RCM and DMS performed RT-qPCR
- RCM performed RT-qPCR analysis
- RCM and DMS performed ICC and imaging
- RCM performed image analysis
- BJP and DMS performed whole genome sequencing on human and chimpanzee PSC lines
- ES assisted with enrichment analyses.
- RCM and AAP designed figures and RCM, DSA, NKS, and DMS generated figures.

## Declaration of generative AI and AI-assisted technologies in the manuscript preparation process

During the preparation of this work, the authors used a large language model to edit portions of the manuscript after writing a complete draft. The authors reviewed and edited the output as needed and take full responsibility for the content of the published article.

